# Age differences in the functional architecture of the human brain

**DOI:** 10.1101/2021.03.31.437922

**Authors:** Roni Setton, Laetitia Mwilambwe-Tshilobo, Manesh Girn, Amber W. Lockrow, Giulia Baracchini, Colleen Hughes, Alexander J. Lowe, Benjamin N. Cassidy, Jian Li, Wen-Ming Luh, Danilo Bzdok, Richard M. Leahy, Tian Ge, Daniel S. Margulies, Bratislav Misic, Boris C. Bernhardt, W. Dale Stevens, Felipe De Brigard, Prantik Kundu, Gary R. Turner, R. Nathan Spreng

**Author notes:** Equal contribution (Roni Setton, Laetitia Mwilambwe-Tshilobo). **Correspondence:** R. Nathan Spreng, Montreal Neurological Institute, 3801 University St., Montréal, QC, H3A 2B4, Canada, p., e.

## Abstract

The intrinsic functional organization of the brain changes into older adulthood. Age differences are observed at multiple spatial scales, from global reductions in modularity and segregation of distributed brain systems, to network-specific patterns of dedifferentiation. Whether dedifferentiation reflects an inevitable, global shift in brain function with age, circumscribed, experience dependent changes, or both, is uncertain. We employed a multi-method strategy to interrogate dedifferentiation at multiple spatial scales. Multi-echo (ME) resting-state fMRI was collected in younger (n=181) and older (n=120) healthy adults. Cortical parcellation sensitive to individual variation was implemented for precision functional mapping of each participant, while preserving group-level parcel and network labels. ME-fMRI processing and gradient mapping identified global and macroscale network differences. Multivariate functional connectivity methods tested for microscale, edge-level differences. Older adults had lower BOLD signal dimensionality, consistent with global network dedifferentiation. Gradients were largely age-invariant. Edge-level analyses revealed discrete, network-specific dedifferentiation patterns in older adults. Visual and somatosensory regions were more integrated within the functional connectome; default and frontoparietal control network regions showed greater connectivity; and the dorsal attention network was more integrated with heteromodal regions. These findings highlight the importance of multi-scale, multi-method approaches to characterize the architecture of functional brain aging.

---

Spontaneous oscillations in brain activity provide the basis for characterizing large-scale functional networks (Biswal et al., 2010; Fox and Raichle, 2007; Yeo et al., 2011). This intrinsic functional network architecture is determined by both genetic factors and experience-dependent neuroplastic changes occurring across timescales, from moments to decades (Stevens and Spreng, 2014). Key organizational features of the intrinsic aging connectome include reduced within- and greater between-network connectivity (Chan et al., 2014; Geerligs et al., 2015), resulting in a dedifferentiated, or less segregated, network architecture (Wig, 2017).

There is now abundant evidence that intrinsic network dedifferentiation is a global feature of functional brain aging (Chan et al., 2014; Geerligs et al., 2015; Stumme et al., 2020 and see Damoiseaux, 2017; Wig, 2017 for reviews). This may reflect functional reorganization in response to systemic structural, neurophysiological or metabolic alterations occurring with age (Reuter-Lorenz and Park, 2014), paralleling the loss of functional specialization within specific brain regions (Cabeza et al., 2002; Park et al., 2004; Rajah and D’Esposito, 2005). These global network changes suggest that network dedifferentiation may be a nonspecific neural marker, paralleling other gross indicators of declining brain health in later life. However, such non-specific changes may also reflect systematic, age-related confounds in data acquisition, processing and analytic approaches to interrogating resting-state fMRI data (Liem et al., 2021 for a review). The most prominent of these is age-related differences in motion (D’Esposito et al., 1999; Geerligs et al., 2015) that introduce spurious age-related differences in connectivity patterns (Power et al., 2012, 2014). These and other confounds highlight the need for rigorous denoising of resting-state fMRI data and pose significant challenges for the field (Power et al., 2018; Spreng et al., 2019).

In contrast to global network changes, there is growing evidence for network-specific patterns of dedifferentiation. Age-related dedifferentiation has been reported among specific association networks (Betzel et al., 2014; Ferriera et al., 2016; Keller et al., 2015; Malagurski et al., 2020; Ng et al., 2016; Rieck et al., 2017; Spreng et al., 2018; Zonneveld et al., 2020) as well as between association and sensorimotor networks (Chan et al., 2014; King et al., 2018; Manza et al., 2020; Meier et al., 2012; Seidler et al., 2015; Song et al., 2014; Stumme et al., 2020). Among the most commonly observed patterns of network-specific dedifferentiation with age is reduced anticorrelation in BOLD signal between the default and dorsal attention networks (Ferreira et al., 2016, Geerligs et al., 2015; Spreng et al., 2016). These canonical brain networks are strongly anticorrelated at rest and during most tasks in younger adults (Fox et al, 2005; Toro et al., 2008; but see Dixon et al., 2017).

Evidence for integration between specific networks hint at a role for resting-state functional connectivity (RSFC) fMRI beyond that of a global indicator of aging brain health. Rather, these specific dedifferentiation patterns may serve as neural markers of domain-specific cognitive changes in later life. Indeed, dissociable patterns of network dedifferentiation have been related to age differences in visuospatial ability (Manza et al., 2020), motor functioning (King et al., 2018), episodic memory (Andrews-Hanna et al., 2007; Chan et al., 2014; Spreng et al., 2018), processing speed (Geerligs et al., 2015; Malagurski et al., 2020; Ng et al., 2016) and executive functioning (Keller et al., 2015; Stumme et al., 2020). Chan and colleagues (2014) provided early evidence that network-specific dedifferentiation patterns were a marker of domain-specific neurocognitive aging. They observed a global pattern of network dedifferentiation (i.e., reduced segregation) that was associated with lower episodic memory ability. However, this relationship was stronger for association networks than for sensorimotor networks, showing that dedifferentiation among cortical association networks was both a sensitive and specific marker of age-related memory decline. We have reported similar findings, showing that age-related increases in connectivity between the default network and frontal regions are associated with lower fluid cognition, as well as specific differences in the nature and content of personal past remembrances in older adults (Spreng et al., 2018).

These reports (and numerous others) suggest that changes to the intrinsic network organization of the brain can provide both global and network-specific markers of neurocognitive aging. However, the range of theoretical, empirical and methodological differences across RSFC studies, as well as numerous analytical challenges, have precluded precise mapping of age-related differences in network organization. Such precision is critically necessary to develop sensitive and specific markers of neurocognitive aging. Here we present an integrated, cross-method study interrogating patterns of network dedifferentiation in a well-powered sample of younger and older adults. We took three approaches to investigate dedifferentiation across multiple spatial scales from global to edgel-level differences, while attempting to mitigate several of the most common analytical challenges. We briefly describe each method and associated hypotheses below. An overview schematic of our analytic approach is presented in Figure 1. Specific methodological details are included in the respective Methods sections.

**Figure 1:**
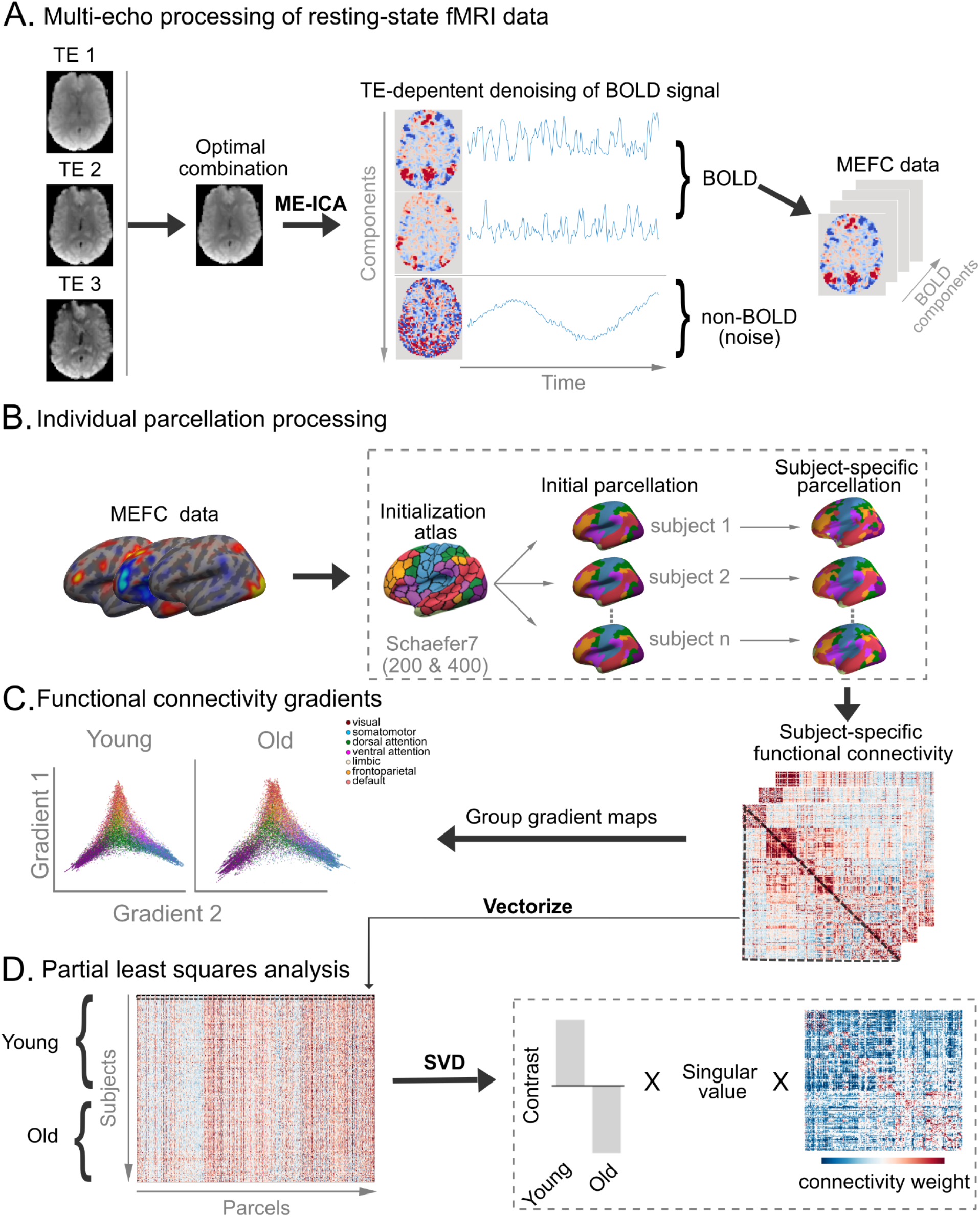
Workflow of study methods. (A) Processing of multi-echo resting-state fMRI images. For each functional run, three echoes (TE_1_, TE_2_, TE_3_) were combined and denoised using multi-echo independent component analysis (ME-ICA). The denoising process involved removing components with non-BOLD signal (noise) and retaining the BOLD components. MEFC images are made up of the BOLD component coefficient sets. (B) Individualized parcellations were generated. The MEFC data for all participants were resampled to a common cortical surface. All participants were first initialized to a pre-defined cortical parcellation atlas (Schaefer atlas). Parcellations were then refined by participant (subject-specific parcellation). For each participant, MEFC data were extracted from and correlated with each parcel to create a subject-specific functional connectivity matrix. These matrices were used to (C) compute cortical gradients in younger and older adults and (D) assess age-related differences in functional connectivity using partial least squares, which performs a singular value decomposition (SVD).

We first examined global network dedifferentiation by measuring differences in spatiotemporal patterns of BOLD signal covariance across the cortex, a measure we refer to as BOLD dimensionality. Innovations in multi-echo fMRI (ME-fMRI) data acquisition protocols, combined with a TE-dependence model of BOLD signal denoising using multi-echo independent components analysis (ME-ICA), enables reliable separation of BOLD from non-BOLD (i.e. noise) signals into different components (Kundu et al., 2017). Emerging evidence suggests that the number of BOLD components, or BOLD dimensionality, is a biologically meaningful metric, showing declines from childhood into middle-age, reflecting greater functional integration and the development of spatially distributed and segregated large-scale networks (Kundu et al., 2018). While this metric has not heretofore been examined in older adulthood, we predicted that BOLD dimensionality, as a proxy for global network dedifferentiation, will be significantly lower for older versus younger adults, consistent with previous reports of age-related network dedifferentiation (Betzel et al., 2014; Chan et al., 2014; Geerligs et al., 2015; Madden et al., 2020; Malagurski et al., 2020; Ng et al., 2016; Stumme et al., 2020; Zonneveld et al., 2019).

Second, we investigated age differences in network dedifferentiation at the macroscale level using a diffusion map embedding approach to estimate RSFC cortical gradients (Huntenburg et al., 2018; Margulies et al., 2016; Paquola et al., 2019; Vos de Wael et al., 2020). Gradient mapping identifies eigenvectors that describe transitions in regional connectivity patterns across the cortical mantle, with a principal RSFC eigenvector solution often differentiating sensory/motor and transmodal cortex (Margulies et al., 2016). As gradients are robust organizational features of the connectome, we predicted that these patterns would be largely resistant to normal age-related changes. However, changes may emerge for regionally-specific connectivity profiles within the macroscale gradient architecture, reflecting network or node specific shifts occurring within an age-invariant macroscale network organization (Bethlehem et al., 2020).

Third, we examined age differences in dedifferentiation patterns identified with unthresholded, edge-level connectomics using Partial Least Squares (PLS) analyses (Krishnan et al., 2011; McIntosh and Misic, 2013). PLS is a multivariate approach that can analyze the full edge-level connectivity matrix in a single statistical step, eliminating the need for additional thresholding within an *a priori* defined network parcellation scheme, enabling us to identify reliable age differences across the full matrix. This provided edge-level precision to detect age differences in the organization of functional brain networks, including patterns of network dedifferentiation. We first examined edge-level connectomics within a canonical seven-network solution (Yeo et al., 2011). We predicted reduced within- and increased between- network connectivity, as reported previously and reviewed above. As we are not aware of any previous studies reporting statistically reliable patterns of unthresholded, edgewise connections, we also anticipated that novel age differences would emerge. Finally, based on previous work (Spreng et al., 2013; Spreng et al., 2016; Grady et al., 2016), we conducted an *a priori* analysis of the sub-network topography for the default, dorsal attention, and frontoparietal control networks, derived from the 17-network solution by Yeo and colleagues (2011). Here we predicted lower within-network connectivity with age, reduced anticorrelation between default and dorsal attention networks, and greater between-network connectivity of the frontoparietal control network with both default and dorsal attention network regions.

Combined, these techniques provide a broad window into the functional architecture of the aging brain, spanning global covariance patterns across the cortex to precision-mapping of edge-level connections. By implementing an integrated, multiscale analytical approach in a well-powered sample of cognitively normal older and younger adults we aimed to characterize both global and specific patterns of RSFC dedifferentiation in the functional architecture of the aging brain.

## Materials and Methods

### Participants

Participants were 181 younger (*M_age_*=22.59y, *SD*=3.27; 57% female) and 120 older (*M_age_*=68.63y, *SD*=6.44; 55% female) healthy adults from Ithaca, New York, and Toronto, Canada (Table 1), rendering a total sample size of 301. Standard inclusion and exclusion criteria were implemented to ensure all participants were healthy without evidence of neurological, psychiatric or other underlying medical conditions known to impact brain or cognitive functioning. Specifically, participants were screened to rule out individuals with acute or chronic psychiatric illness. Participants were also queried for current usage of medications for mood (e.g. depression), thinking or mental abilities (e.g. attention deficit disorder) or having experienced significant changes to health status within three months of the eligibility interview. Younger and older participants were screened for depressive symptoms using the Beck Depression Inventory (Beck et al., 1996) or the Geriatric Depression Scale (Yesavage et al., 1982), respectively. Two older adults were excluded due to a rating of “moderate depression”. In order to screen for normal cognitive functioning, participants were administered the Mini-Mental State Examination (MMSE; Folstein et al., 1975; M_younger_: 29.1; SD_younger_: 1.2; M_older_: 28.6; SD_older_: 1.3) and NIH cognition battery (Gershon et al., 2013). If participants performed below 27/30 on MMSE and scored in the bottom 25th percentile of age-adjusted scores for fluid cognition index on the NIH (Hacket et al., 2018; Scott et al., 2019), they were excluded. All participants were right-handed with normal or corrected-to-normal vision. Procedures were administered in compliance with the Institutional Review Board at Cornell University and the Research Ethics Board at York University.

**Table 1:** Sample Demographics. Episodic Memory, Semantic Memory, and Executive Function are index scores. Processing Speed is a z-score on Symbol Digit Modalities Task, Oral. * significant group differences. Education was not recorded for 14 participants. Age group differences in episodic memory, semantic memory, executive function, and processing speed were tested in 283 participants. Positive T values reflect higher scores in younger adults, negative values reflect higher scores in older adults. Statistical results were nearly identical when including sex, education, site, and estimated whole brain volume as covariates in an ANCOVA.

|  | Descriptive Statistics |  | Inferential Statistics |  |  |  |  |
| --- | --- | --- | --- | --- | --- | --- | --- |
|  | Younger Adults | Older Adults | *T* | dof | *p* | 95% CI | Cohen's *d* |
| N |  |  |  |  |  |  |  |
| Cornell | 154 (86 female) | 84 (47 female) |  |  |  |  |  |
| York | 27 (17 female) | 36 (19 female) |  |  |  |  |  |
| Race | 60.38% white,<br>19.50% asian,<br>8.18% black,<br>5.03% other,<br>4.41% mixed,<br>2.50% not provided | 92.38% white,<br>2.54% asian,<br>2.54% black,<br>2.54% other |  |  |  |  |  |
| Ethnicity | 81.76% non-hispanic or latino,<br>10.69% hispanic or latino,<br>7.55% not provided | 89.83% non-hispanic or latino,<br>8.48% not provided,<br>1.69% hispanic or latino, |  |  |  |  |  |
| Age (years) |  |  |  |  |  |  |  |
| Range | 18-34 | 60-89 |  |  |  |  |  |
| *M* | 22.6 | 68.6 |  |  |  |  |  |
| *SD* | 3.3 | 6.4 |  |  |  |  |  |
| Education (years)* |  |  | -7.20 | 285 | < .001 | [-2.5, -1.48] | 0.86 |
| Range | 12-24 | 12-24 |  |  |  |  |  |
| *M* | 15.2 | 17.2 |  |  |  |  |  |
| *SD* | 1.9 | 2.9 |  |  |  |  |  |
| Episodic Memory* |  |  | 17.51 | 281 | < .001 | [1.1, 1.38] | 2.11 |
| Range | -1.75-1.59 | -1.99-0.70 |  |  |  |  |  |
| *M* | 0.52 | -0.71 |  |  |  |  |  |
| *SD* | 0.53 | 0.66 |  |  |  |  |  |
| Semantic Memory* |  |  | -9.18 | 281 | < .001 | [-1.00, -.65] | 1.10 |
| Range | -2.78-1.39 | -1.29-1.91 |  |  |  |  |  |
| *M* | -0.35 | 0.48 |  |  |  |  |  |
| *SD* | 0.77 | 0.71 |  |  |  |  |  |
| Executive Function* |  |  | 12.67 | 281 | < .001 | [.71, .97] | 1.52 |
| Range | -1.15-1.80 | -2.03-0.76 |  |  |  |  |  |
| *M* | 0.36 | -0.48 |  |  |  |  |  |
| *SD* | 0.56 | 0.53 |  |  |  |  |  |
| Processing Speed* |  |  | 15.03 | 281 | < .001 | [1.17, 1.53] | 1.81 |
| Range | -2.26-3.05 | -2.40-.050 |  |  |  |  |  |
| *M* | 0.57 | -0.78 |  |  |  |  |  |
| *SD* | 0.86 | 0.56 |  |  |  |  |  |

### Cognitive Assessment

We first characterized our sample with a deep cognitive assessment. 283 of 301 individuals (163/181 younger adults, 120/120 older adults) underwent cognitive testing prior to brain scanning. Index scores were created for cognitive domains of episodic memory, semantic memory, executive function, and processing speed (descriptives in Table 1). Episodic memory tasks included Verbal Paired Associates from the Wechsler Memory Scale-IV (Wechsler, 2009), the Associative Recall Paradigm (Brainerd et al., 2014), and NIH Cognition Toolbox Rey Auditory Verbal Learning and Picture Sequence Memory Tests (Gershon et al., 2013). Semantic memory tasks included Shipley-2 Vocabulary (Shipley et al., 2009), and NIH Cognition Toolbox Picture Vocabulary and Oral Reading Recognition Tests (Gershon et al., 2013). Executive function comprised the Trail Making Test (B-A; Reitan, 1958), the Reading Span Task (Daneman & Carpenter, 1980), NIH Cognition Toolbox Flanker Inhibitory Control and Attention task, Dimensional Change Card Sort, and List Sort Working Memory Tests (Gershon et al., 2013). Processing speed was tested with the Symbol Digit Modalities Test, Oral (Smith, 1982).

All data were z-scored. Index scores represent the average z-score for all measures included within a cognitive domain. Across the four domains, higher scores represent better performance. Brain-behavior product-moment correlations were conducted at an alpha level of .05 with 95% confidence intervals. Bonferroni adjustments for multiple comparisons were set at *p* < .013 for the four index score tests.

### Neuroimaging

#### Image Acquisition

Imaging data were acquired on a 3T GE750 Discovery series MRI scanner with a 32-channel head coil at the Cornell Magnetic Resonance Imaging Facility or on a 3T Siemens Tim Trio MRI scanner with a 32-channel head coil at the York University Neuroimaging Center in Toronto. Scanning protocols were closely matched across sites. Anatomical scans at Cornell were acquired using a T1-weighted volumetric magnetization prepared rapid gradient echo sequence (TR=2530ms; TE=3.4ms; 7° flip angle; 1mm isotropic voxels, 176 slices, 5m25s) with 2x acceleration with sensitivity encoding. At York, anatomical scans were acquired using a T1-weighted volumetric magnetization prepared rapid gradient echo sequence (TR=1900ms; TE=2.52ms; 9° flip angle; 1mm isotropic voxels, 192 slices, 4m26s) with 2x acceleration and generalized auto calibrating partially parallel acquisition (GRAPPA) encoding at an iPAT acceleration factor of 2. Two 10m06s resting-state runs were acquired using a multi-echo (ME) EPI sequence at Cornell University (TR=3000ms; TE_1_=13.7ms, TE_2_=30ms, TE_3_=47ms; 83° flip angle; matrix size=72×72; field of view (FOV)=210mm; 46 axial slices; 3mm isotropic voxels; 204 volumes, 2.5x acceleration with sensitivity encoding) and York University (TR=3000ms; TE_1_=14ms, TE_2_=29.96ms, TE_3_=45.92ms; 83° flip angle; matrix size=64×64; FOV=216mm; 43 axial slices; 3.4×3.4×3mm voxels; 200 volumes, 3x acceleration and GRAPPA encoding). Participants were instructed to stay awake and lie still with their eyes open, breathing and blinking normally in the darkened scanner bay.

#### Image Processing

Anatomical images were skull stripped using the default parameters in FSL BET (Smith, 2002). Brain-extracted anatomical and functional images were submitted to ME independent component analysis (ME-ICA; version 3.2 beta; https://github.com/ME-ICA/me-ica; Kundu et al., 2011; Kundu et al., 2013). ME-ICA relies on the TE-dependence model of BOLD signal to determine T2* in every voxel and separates BOLD signal from non-BOLD sources of noise. Prior to TE-dependent denoising, time series data were minimally preprocessed: the first 4 volumes were discarded, images were computed for de-obliquing, motion correction, and anatomical-functional coregistration, and volumes were brought into spatial alignment across TEs. The T2* maps were then used for anatomical-functional coregistration. Grey matter and cerebrospinal fluid compartments are more precisely delineated by the T2* map than by raw EPI images (Speck et al., 2001; Kundu et al., 2017), which is an important consideration in aging research where these boundaries are often blurred by enlarged ventricles and greater subarachnoid space. Volumes were then optimally combined across TEs and denoised. The outputs of interest included: 1) spatial maps consisting of the BOLD components, 2) reconstructed time series containing only BOLD components, and 3) the BOLD component coefficient sets.

Image quality assessment was performed on the denoised time series in native space to identify and exclude participants with unsuccessful coregistration, residual noise (framewise displacement > .50 mm coupled with denoised time series showing DVARS >1, Power et al., 2012), temporal signal to noise ratio < 50, or fewer than 10 retained BOLD-like components (see Supplementary Figure 1 for the group temporal signal to noise map).

The denoised BOLD component coefficient sets in native space, optimized for functional connectivity analyses (Kundu et al., 2013), were used in subsequent steps. We refer to these as multi-echo functional connectivity (MEFC) data. Additional measures were taken to account for variation in the number of independent components from ME-ICA once connectivity matrices were estimated, as detailed below. MEFC neuroimages were mapped to a common cortical surface for each participant using FreeSurfer v6.0.1 (Fischl et al., 2012). To maximize alignment between intensity gradients of structural and functional data (Greve & Fischl, 2009), MEFC data were first linearly registered to the T1-weighted image by run. The inverse of this registration was used to project the T1-weighted image to native space and resample the MEFC data onto a cortical surface (fsaverage5) with trilinear volume-to-surface interpolation. This produces a cortical surface map where each vertex, or surface point, is interpolated from the voxel data. Once on the surface, runs were concatenated and MEFC data at each vertex were normalized to zero mean and unit variance.

##### Individualized Parcellation

ME-fMRI processed data provides excellent reliability and temporal signal-to-noise, sufficient for individual-subject precision mapping (Lynch et al., 2020; Lynch, Elbau, and Liston, 2021). An individualized functional parcellation approach was implemented to identify person-specific functional network nodes (Chong et al., 2017). These individualized parcellations were used in both the gradient and edge-level connectivity analyses to facilitate comparisons of RSFC between younger and older adults. Poor registration to standardized templates may fail to capture individual variability in functional organization of the cortex, and these registration problems may systematically differ across age groups (Braga and Buckner, 2017; Chong et al., 2017; Gordon, et al. 2017; Kong et al., 2019; Kong et al., 2021; Laumann et al., 2015; Wang et al., 2015). Deriving functionally-defined, person-specific cortical parcellations can account for differences at the level of the individual, thereby mitigating systematic registration biases in between-group comparisons. Adopting an individualized parcellation approach may also lessen the impact of noise artifacts that can obscure small yet reliable group differences, increasing power to detect reliable brain-behavior associations (Kong et al., 2021).

We generated participant-specific functional parcellations to examine individual differences in functional brain network organization using the Group Prior Individual Parcellation (GPIP; Chong et al., 2017). This approach enables a more accurate estimation of participant-specific individual functional areas (Chong et al., 2017) and is more sensitive to detecting RSFC associations with behavior (e.g. Kong et al., 2019; Mwilambwe-Tshilobo et al., 2019). The main advantage of this approach is that the correspondence among parcel labels is preserved across participants, while the parcel boundaries are allowed to shift based on the individual-specific functional network organization of each participant—thus providing a similar connectivity pattern that is shared across the population. Starting from an initial pre-defined group parcellation atlas, GPIP first refines each individual’s parcel boundaries relative to their resting-state fMRI data. Next, the concentration (inverse covariance/partial correlation) matrices from all subjects are jointly estimated using a group sparsity constraint. GPIP iterates between these two steps to continuously update the parcel labels until convergence, defined as no more than one vertex changing per parcel or 40 iterations. Compared to other group-based parcellation approaches, GPIP has shown to improve the homogeneity of the BOLD signal within parcels and the delineation between regions of functional specialization (Chong et al., 2017).

We extracted MEFC data from each vertex and applied the above parcellation across the entire cohort of 301 participants at resolutions of 200 and 400 parcels. For each resolution, MEFC data were initialized to a group parcellation atlas developed by Schaefer et al. (2018). We use this cortical parcellation scheme both as the initialization reference for our individualized cortical parcellation maps, as well as for our edge-level connectomic analyses described below. We selected the Schaefer atlas for three reasons: (i) It is functionally-derived, and thus more closely aligned with the current study aims, (ii) it has high spatial resolution for different levels of granularity (we report findings from both 400 and 200 node parcellations here), and (iii) it is among the most commonly reported cortical parcellation atlases in the literature, providing intrinsic partitioning of nodes within Yeo 7- and 17- network solutions (used in our edge-level connectomics analyses; Yeo et al., 2011).

Following initialization with the Schaefer parcellations, the two-step iterative process was repeated 20 times to produce a final parcellation representing the optimal partition with respect to the entire cortical surface. We calculated homogeneity by taking the average correlation coefficient of all pairs of vertices in a given parcel and then averaging across all parcels. This was repeated at each repetition to observe the incremental change in homogeneity as the iterative parcellation proceeded. Homogeneity was calculated first at the participant level and then averaged across the entire cohort for a group estimate. For a subset of participants, some parcels from the final partition merged into the medial wall (where no data existed) or into parcels belonging to the contralateral hemisphere. Because partitions likely reflect participant-specific neurobiological variations in functional organization, parcels assigned to the contralateral hemisphere were allowed to retain their original group atlas labels. With the 400-parcel resolution, parcels merging with the medial wall occurred in 69 older adults and 35 younger adults, averaging 2-3 parcels in these participants; parcels migrating to the contralateral hemisphere occurred in 62 older adults and 24 younger adults, averaging 2-3 parcels. With the 200-parcel resolution, parcels merging with the medial wall occurred in 18 older adults and 10 younger adults, averaging 1 parcel in these participants. No parcels migrated to the contralateral hemisphere at this resolution.

##### Functional Connectivity Matrix

A connectivity matrix was constructed for each participant according to their individualized parcel solution. Since the MEFC data consist of ICA coefficient sets (coefficient weights for each accepted component x vertex) concatenated by run, we extracted and averaged the MEFC data from vertices within each parcel to obtain a parcel-level coefficient set. Connectivity was estimated by computing the product-moment correlation between each parcel’s coefficient set, resulting in a n_parcels_ × n_parcels_ functional connectivity matrix (Ge, Holmes, Buckner, Smoller, & Sabuncu, 2017). In this approach, RSFC was calculated as the correlation of the ICA coefficients across parcels, rather than a correlation across BOLD signal time-series, as is typically done (see Kundu et al., 2013). The canonical Fisher’s r-to-z transformation was then applied to normalize the distribution of correlation values and account for variation in MEFC data degrees of freedom, or the number of denoised ICA coefficients (i.e. number of BOLD components), across individuals (Kundu et al., 2013):

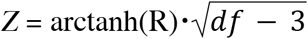

where R is the product-moment correlation value and *df* is the number of denoised ICA coefficients. Computing functional connectivity with approximately independent component coefficients rendered global signal regression unnecessary (Spreng et al., 2019). Critically, ME-ICA effectively removes distance-dependent RSFC motion confounds from fMRI data (Power et al., 2018). As shown in Supplemental Figure 2 (see also Supplemental Material), framewise displacement had a comparable impact on younger and older adult RSFC, ruling out motion as a potential confound in the results reported below.

### Analysis

#### BOLD Dimensionality

A unique advantage of ME-fMRI and the ME-ICA processing framework is that BOLD- and non-BOLD-like signals are separated into independent components. A novel metric of “BOLD dimensionality,” the number of BOLD components identified by ME-ICA, may then be examined as a data-driven representation of the global network architecture of the brain, and used to investigate changes with age (e.g. Kundu et al., 2018). We assessed the test-retest reliability of BOLD dimensionality across two runs of data. Total BOLD dimensionality was then compared between age-groups with an independent samples *t*-test and an ANCOVA controlling for sex, education, site and estimated whole brain volume (eWBV; sum of grey and white matter divided by total intracranial volume, derived from FreeSurfer). To observe the trajectory of BOLD dimensionality with increasing age across the lifespan, BOLD dimensionality data from an independent developmental sample (N = 51, 10 female; *M_age_*=21.9 years; age range, 8.3 – 46.2 years; see Kundu et al., 2018 for details) were pooled with the current data. As our sample consisted of two discrete age cohorts, these additional data points were used to properly fit a function between age and BOLD dimensionality. To render the samples comparable and account for differences in acquisition across datasets, BOLD dimensionality was scaled by the number of timepoints acquired. The relationship between age and BOLD dimensionality was then fit to a power law function (see Supplemental Figure 3 for unscaled version). Further characterization of BOLD signal dimensionality, including associations with graph analytic measures of participation coefficient, modularity and segregation, and BOLD signal dimensionality’s relationship to whole brain RSFC are reported in Supplemental Material (Supplemental Table 1, Supplemental Figures 4 & 5)

#### Gradients & Manifold Eccentricity

Cortical gradients allow for a low dimensional (i.e. macroscle) representation of functional connectivity that demarcate transitions in whole brain functional connectivity (Margulies et al., 2016; Huntenburg et al. 2018). Gradient map embedding has reliably demarcated cortical transitions from unimodal to heteromodal cortex, visual to somatomotor cortices, among others (Bethlehem et al. 2020; Margulies et al., 2016; Hong et al. 2020). Cortical gradients were computed using functions from the BrainSpace toolbox (https://github.com/MICA-MNI/BrainSpace; Vos de Wael et al., 2020), as implemented in MATLAB. For each participant, the 400 x 400 GPIP functional connectivity matrix was thresholded row-wise to the upper 10% of connections to retain only the strongest positive connections (Hong et al., 2019; Margulies et al., 2016). Cosine similarity was computed on the sparse matrix to input to the diffusion map embedding algorithm employed below, generating a matrix that captures similarity in whole-brain connectivity patterns between vertices (Hong et al., 2019; Margulies et al., 2016).

We then applied diffusion map embedding, a non-linear dimensionality manifold learning technique from the family of graph Laplacians (Coifman et al., 2005), to identify gradient components at the individual participant level. Each gradient represents a low-dimensional embedding/eigenvector estimated from a high-dimensional similarity matrix. In the embedding space, vertices that feature greater similarity in their whole-brain functional connectivity patterns appear closer together, whereas vertices that are dissimilar are farther apart. Each embedding axis can thus be interpreted as an axis of variance based on connectivity pattern similarity/dissimilarity. Euclidean distance in the embedded space is equivalent to the diffusion distance between probability distributions centered at those points, each of which is equivalent to a *difference in gradient* score. The algorithm is controlled by a single parameter α, which controls the influence of density of sampling points on the manifold (Margulies et al, 2016). We used α = 0.5 in this study, which differentiates diffusion map embedding from Laplacian eigenmaps, and allows the inclusion of both global and local relationships in the estimation of the embedded space. An iterative Procrustes rotation was performed to align participant-specific gradient components to a young-old group average template and enable group comparisons. Group contrasts were conducted using surface-based linear models, as implemented in Surfstat (Worsley et al., 2009; http://www.math.mcgill.ca/keith/surfstat/) controlling for sex, education, site and eWBV.

We calculated a metric of manifold eccentricity to quantify the diffusivity of vertices in gradient space. Following Bethlehem et al. (2020) and Park et al. (2021), we summed the squared Euclidean distance of each vertex from the whole-brain median in a 2-dimensional gradient space for each participant. The position of a vertex in gradient space represents a coordinate for where the vertex falls on each gradient’s axis. The proximity of any two vertices informs how similar their functional connectivity profiles are on each gradient. The more diffuse vertices are within a network, the more variable and dedifferentiated the functional connectivity profiles. Mean manifold eccentricity was then compared across age groups. Statistical significance was determined with non-parametric spin-test permutation testing, which overcomes biases in the test statistic due to the spatial autocorrelation inherent to BOLD data (Alexander-Bloch et al., 2018). An ANCOVA on manifold eccentricity was also conducted controlling for sex, education, site and eWBV.

#### Edge-Level Connectomics

Inter-regional functional connectivity group differences were tested with PLS. PLS is a multivariate method that determines the association between two sets of variables by identifying linear combinations of variables in both sets that maximally covary together (McIntosh and Lobaugh, 2004; McIntosh and Misic, 2013). Crucially, PLS enables whole-brain contrasts of unthresholded connectivity matrices, allowing more precise mapping of edgewise age differences. In our analyses, one set of variables was individual RSFC matrices, while the other set represented group assignment or individual difference metrics (e.g., BOLD dimensionality; see Supplemental Material).

Functional connectivity was assessed at the whole-brain level using the Schaefer atlas (Schaefer et al., 2018; Yeo et al., 2011; 400 x 400 matrix; 200 x 200 matrix as supplementary analysis, Supplemental Figure 6). Motivated by prior work (e.g., Grady et al., 2016; Sullivan et al., 2019; Spreng et al., 2016), we also examined RSFC among sub-networks of the default, dorsal attention, and frontoparietal control networks. For the sub-network analysis, we first reassigned each of the 400 parcels to the corresponding network of the Yeo 17-network solution following the mapping based on Schaefer et al. (2018). Next, we created a matrix for the pairwise connections between 8 sub-networks: dorsal attention (DAN-A, DAN-B), frontoparietal control (CONT-A, CONT-B, CONT-C), and default (DN-A, DN-B, DN-C), resulting in a 192×192 parcel matrix. The full 17-network characterization of the 400×400 parcel results, along with the 17-network and sub-network characterizations of the 200×200 matrix, can be found in Supplemental Figures 7, 8, and 9. At each level, a data matrix **X** was created using all participants’ parcellated functional connectivity matrices. The **X** matrix was organized such that each row corresponded to an observation (each participant, nested in age groups), and the cells in each column corresponded to the unique connections from each participant’s connectivity matrix (the lower triangle of the matrix). The column means within each group were calculated, and the data in **X** were mean-centered. The mean-centered data were then submitted to singular value decomposition (SVD) to provide mutually orthogonal latent variables. Each latent variable represents a specific relationship (e.g. RSFC x Group) and consists of three elements: (1) a left singular vector consisting of the weighted connectivity pattern that optimally expresses the covariance, (2) a right singular vector, which represents the weights of the study design variables and can be interpreted as data-driven contrast weights between groups, and (3) a scalar singular value, which represents the covariance strength between the design variables (Group) and RSFC accounted for by each latent variable. Brain connectivity scores were calculated by taking the dot product of the left singular vector and each participant’s RSFC matrix. A brain connectivity score, therefore, represents a single measure of the degree to which a participant expresses the connectivity pattern captured by a given latent variable.

All PLS latent variables were statistically evaluated using permutation testing. Rows of **X** were randomly reordered and subjected to SVD iteratively, as described above. This was done 1,000 times to create a distribution of singular values under the null hypothesis of no existing relationships between X and Y for the corresponding PLS analysis: that there is no group difference in whole-brain (or sub-network) RSFC. A *p*-value was computed for each latent variable as the proportion of permuted singular values greater than or equal to the original singular value. Critically, permutation tests involve the entire multivariate pattern and are performed in a single analytic step, so correction for multiple comparisons is not required (McIntosh and Lobaugh, 2004).

Bootstrap resampling was used to estimate the reliability of weights for each RSFC edge. Participants were randomly resampled (rows in **X**) with replacement while respecting group membership. The matrix was subjected to SVD and the process was repeated 1,000 times, generating a sampling distribution for the weights in the singular vectors. To identify individual connections that made a statistically significant contribution to the overall connectivity pattern, we calculated the ratio between each weight in the singular vector and its bootstrap-estimated standard error. Bootstrap ratios are equivalent to z-scores if the bootstrap distribution is approximately unit normal (Efron and Tibshirani, 1986). Bootstrap ratios were, therefore, thresholded at values of ±1.96, corresponding to the 95% CI.

#### Network-Level Contributions

PLS analyses identified inter-regional connectivity patterns that differed by group and/or covaried with individual difference metrics. For each of these analyses, network-level effects were also examined. To quantify the network-level contributions to the PLS-derived functional connectivity pattern, two separate weighted adjacency matrices were constructed from positive and negative RSFC weights. For both matrices, nodes represent parcels defined by the individual parcellation, while edges correspond to the thresholded bootstrap ratio of each pairwise connection. Network-level functional connectivity contributions were quantified by assigning each parcel according to the network assignment reported by Yeo et al. (2011), and taking the average of all connection weights in a given network, thereby generating a 7 x 7 matrix (17 x 17 matrix for the 17-network solution; and an 8 x 8 matrix when examining the default, dorsal attention, and frontoparietal control sub-networks). The significance of mean within- and between- network connectivity was computed by permutation testing. During each permutation, network labels for each node were randomly reordered and the mean within- and between-network connectivity were recalculated. This process was repeated 1000 times to generate an empirical null sampling distribution that indicates no relationship between network assignment and connectivity pattern (Shafiei et al., 2019). The significance of the pairwise connections to the network matrix was determined by estimating the proportion of times the value of the sampling distribution was greater than or equal to the original value.

#### Spring-Embedded Plots

Spring-embedded plots were rendered from group average matrices of RSFC data using Pajek software (Mrvar and Batagelj, 2016). Sparse matrices containing the top 5% of positive connections were entered into Pajek. The plotting of positive edge weights and similar thresholds applied to prior investigations of healthy aging (Chan et al., 2014; Geerligs et al., 2015) permit more direct comparison with prior results. A partition was assigned based on the Yeo 7- or 17-network solution (Yeo et al., 2011) to optimize community (i.e., network) structure for visualization.

## Results

To interrogate the intrinsic functional architecture of the aging brain, we implemented a multifaceted, multiscale data acquisition and analysis protocol in younger and older healthy adults (see Figure 1 and Methods). To identify global patterns of network dedifferentiation with age, we first assessed age differences in the dimensionality of the ME-fMRI BOLD signal as output from ME-ICA. Next, we examined network-specific dedifferentiation patterns, contrasting macroscale gradients and edge-level network connectomics between younger and older adults. At each turn, we examined associations between network organization and cognitive functioning for younger and older adults. Brain and behavior associations for each analysis are reported in Supplemental Materials (Supplemental Tables 2, 3 and 4; Supplemental Figures 10 and 12). All results are reported with covariates of site, sex, education, and eWBV where appropriate.

### BOLD Dimensionality

Two 10-minute runs of resting-state ME-fMRI were collected. BOLD dimensionality, the number of independent BOLD components in ME-fMRI signal, was stable across runs (*r*(299) = .79, *p* < .001 [.75, .83]; Figure 2A). Younger adults showed greater BOLD dimensionality than older adults (*t*(299)=15.38, *p* < .001; Cohen’s *d*= 1.81; Figure 2B). This remained true when covariates of site, sex, education, and eWBV were included (*F*(1,281)= 97.07, *p* < .001; η_p_ = .26). In the context of lifespan development, which included an additional sample aged 8-46 (Kundu et al., 2018), a power function provided a suitable fit between age and BOLD dimensionality (R^2^=.547; Figure 2C). BOLD dimensionality associations with cognition are in Supplemental Tables 2, 3 and 4 and Supplemental Figure 10A.

**Figure 2:**
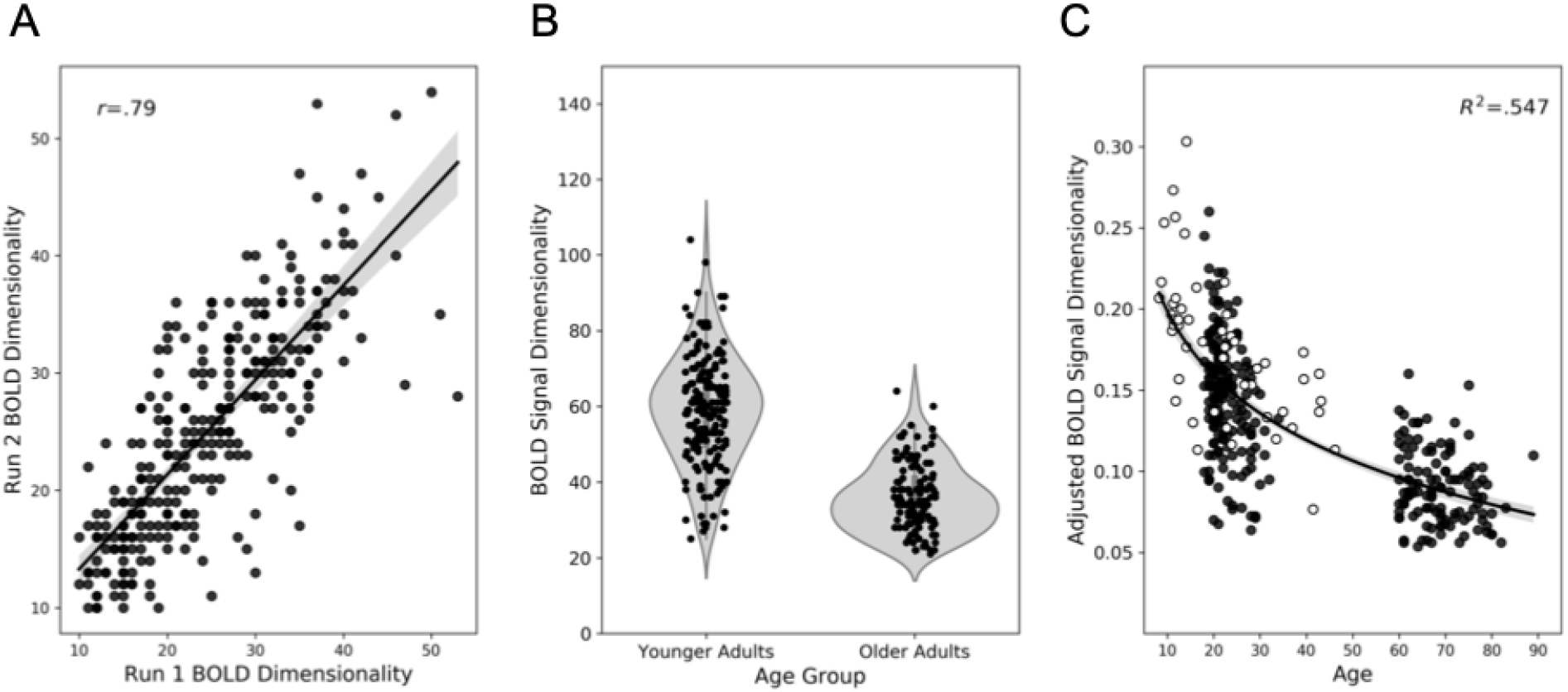
BOLD signal dimensionality. (A) High test-retest reliability across two ME-fMRI runs. (B) Violin plots show distributions of total BOLD signal dimensionality across runs in younger and older adults. (C) Scatter plot showing BOLD signal dimensionality by age with a power distribution and 95% confidence intervals overlaid. Points in white were contributed by Kundu and colleagues (2018). Adjusted BOLD signal dimensionality = Total number of accepted BOLD components / number of time points acquired.

### Gradient Analyses

We next characterized macroscale gradients of RSFC (e.g., Hong et al., 2019; Margulies et al., 2016) in younger and older adults. In both groups, the principal gradient ran from sensory and motor regions towards transmodal systems such as the default network (Figure 3A), suggesting that macroscale functional organization of the cortex is generally preserved with age. However, regional age differences in this topographic organization emerged (FWE *p* < .05; cluster defining threshold *p* < .01; Figure 3A). Indeed, cortex-wide age group comparison on the principal gradient revealed higher gradient values in the right superior parietal lobule and somatosensory cortex, but lower values in occipital and ventral temporal regions for older adults.

**Figure 3:**
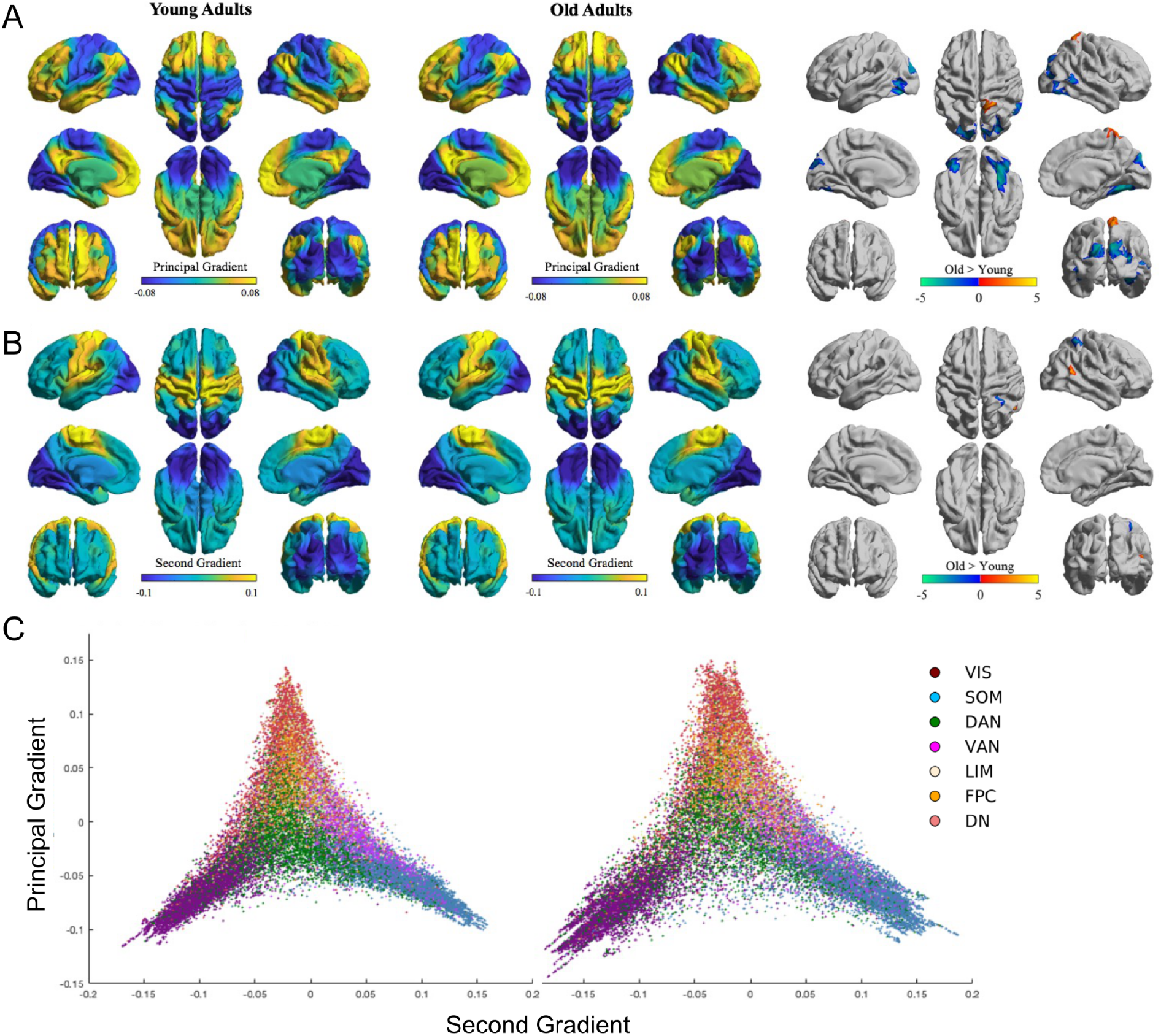
Gradients of cortical connectivity in younger and older adults. (A) The mean principal gradient for younger (left) and older (center) adults, representing an axis of functional connectivity similarity variance that ranged lowest to highest from unimodal to transmodal cortex. (B) The mean second gradient for younger (left) and older (center) adults, representing an axis of functional connectivity similarity variance that ranged lowest to highest from visual to somatomotor cortex. Older adults > younger adults contrasts revealing statistically significant clusters at FWE *p* < 0.05, cluster defining threshold *p* < 0.01 (A & B right). (C) Vertex-wise scatterplots representing the principal-second gradient values. This gradient manifold is depicted for younger (left) and older (right) adults. Scatterplot colors indicate functional networks as per the 7-network solution by Yeo et al. (2011). VIS = visual, SOM= somatomotor, DAN= dorsal attention, VAN = ventral attention, LIM = limbic, FPC= frontoparietal control, DN= default.

A difference between visual and sensory/motor networks with respect to cortical gradient organization was also evident when examining the second gradient. As in prior studies (e.g. Margulies et al., 2016), the second gradient differentiated visual from somatomotor cortices in both groups (Figure 3B). However, cortex-wide between-group comparisons revealed subtle differences in this topography. In particular, we observed increased gradient values in the temporoparietal junction in older compared to younger adults, together with decreased values in a segment of the superior parietal lobule/intraparietal sulcus. These results show that regions along the second gradient axis also shift their connectivity profiles with advancing age, again with shifts in sensory/visual regions.

Finally, we rendered principal-second gradient manifold scatterplots in a 2D gradient embedding space in younger and older adults (Figure 3C). Older adults showed more diffuse, and, thus, dedifferentiated vertices. We quantified this diffusivity by calculating manifold eccentricity– the sum of Euclidean distance across all vertices from the median– for each participant and compared across groups. Results revealed significantly greater manifold eccentricity in older adults (*t*(299) = −10.74, *p*_SPIN_ < 0.01, Cohen’s *d* = 1.26; *F*(1,281)= 47.18, *p* < .001, η_p_^2^ = .14 with site, sex, education, and eWBV covariates included). See Supplemental Tables 2, 3 and 4 and Supplemental Figure 10B for associations with behavior.

As BOLD dimensionality and manifold eccentricity both demonstrated significant age group differences, we conducted post-hoc product-moment correlations to test whether these global measures of brain organization were reliably associated. Negative correlations were observed in both younger (*r*(179)= -.575, *p* < .001, [-.66, -.47]) and older adults (*r*(118)= -.255, *p* < .005, [-.42, -.08]), such that higher BOLD dimensionality was related to less diffuse, more compact vertices in the manifold. In computing a partial correlation controlling for age, the relationship remained when performed on the full sample (*pr*(298)= -.391, *p* < .001, [-.48, -.29]). Non-overlapping 95% confidence intervals indicated a significantly more negative correlation in younger adults. Results were similar when repeated with covariates (young: *pr*(161)= -.45, *p* < .001, [-.53, -.36]; old: *pr*(113)= -.23, *p* < .05, [-.35, -.11]; full sample: *pr*(280)= -.34, *p* < .001, [-.44, -.23]), although confidence intervals overlapped between groups. Supplemental Figure 11 illustrates the relationship in each age group.

### Edge-Level Connectomics

We next examined edge-level, interregional functional connectivity differences between younger and older adults. Group mean connectivity matrices are in Figure 4A-B. Qualitative differences in the top 5% of positive connections between groups can be observed with a spring-embedded layout arranged by network membership (Figure 4C-D). The spring-embedded plot suggests more integration of the dorsal attention and frontoparietal control networks in older adults.

**Figure 4:**
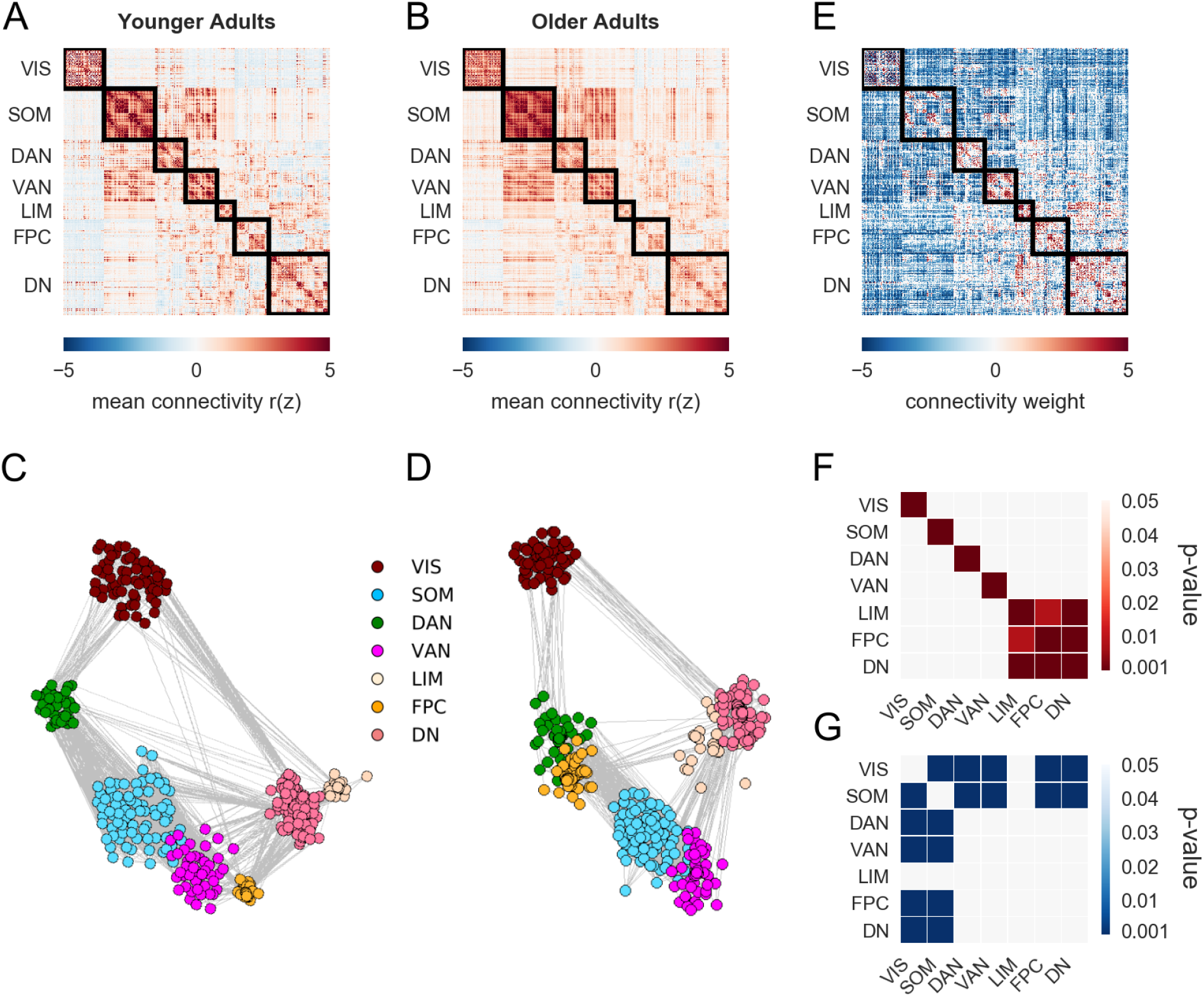
Functional connectomics in younger and older adults. Mean RSFC for the 400-parcellated data in (A) younger and (B) older adults. Spring-embedded plots with a 7-network solution (5% edge density) of the mean correlation matrices for (C) younger and (D) older adults. Nodes that are highly functionally correlated with one another are grouped closer together. (E) Multivariate PLS analysis was used to identify age-related differences in RSFC between younger and older adults. Red color indicates significantly greater RSFC in younger adults, and blue color indicates significantly greater RSFC in older adults. (F-G) Network contributions represent the summary of significant positive and negative edge weights within and between networks in younger (F) and older (G) adults. The mean positive and negative bootstrap ratios within and between networks are expressed as a *p*-value for each z-score relative to a permuted null model. Higher z-scores indicate greater connectivity than predicted by the null distribution. VIS = visual, SOM = somatomotor, DAN = dorsal attention, VAN = ventral attention, LIM = limbic, FPC = frontoparietal control, DN = default.

#### PLS (whole brain)

Age-related differences in the 79800 interregional connections (i.e., the lower triangle of the 400×400 functional connectivity matrix) were quantitatively assessed with PLS. A significant latent variable (permuted *p* < .001) revealed a pattern of age differences in RSFC, with increases and decreases observed across the connectome (Figure 4E). Network contribution analysis of within- and between- network edges revealed significant age effects. Older adults demonstrated lower within-network connectivity across all seven networks, and lower connectivity between limbic, frontoparietal control and default networks (Figure 4F). Older adults showed greater between-network connectivity across systems for the visual and somatomotor networks (Figure 4G). The overall pattern of age-related differences was similar when examined with a 200 parcellation scheme (Supplementary Figure 6). Brain connectivity scores’ association with cognition are reported in Supplemental Table 2, 3, and 4, and Supplemental Figure 12.

#### PLS (sub-network)

In an *a priori*, targeted sub-network analysis we examined age-group differences in functional connectivity among sub-networks of the default, dorsal attention, frontoparietal control networks. The mean age-group sub-network matrices are shown in Figure 5A-B. The spring-embedded representation of the top 5% of positive connections in each group (Figure 5C-D) suggests that older adults show more integration of the default network (DN-A) and frontoparietal control network (CONT-C).

**Figure 5:**
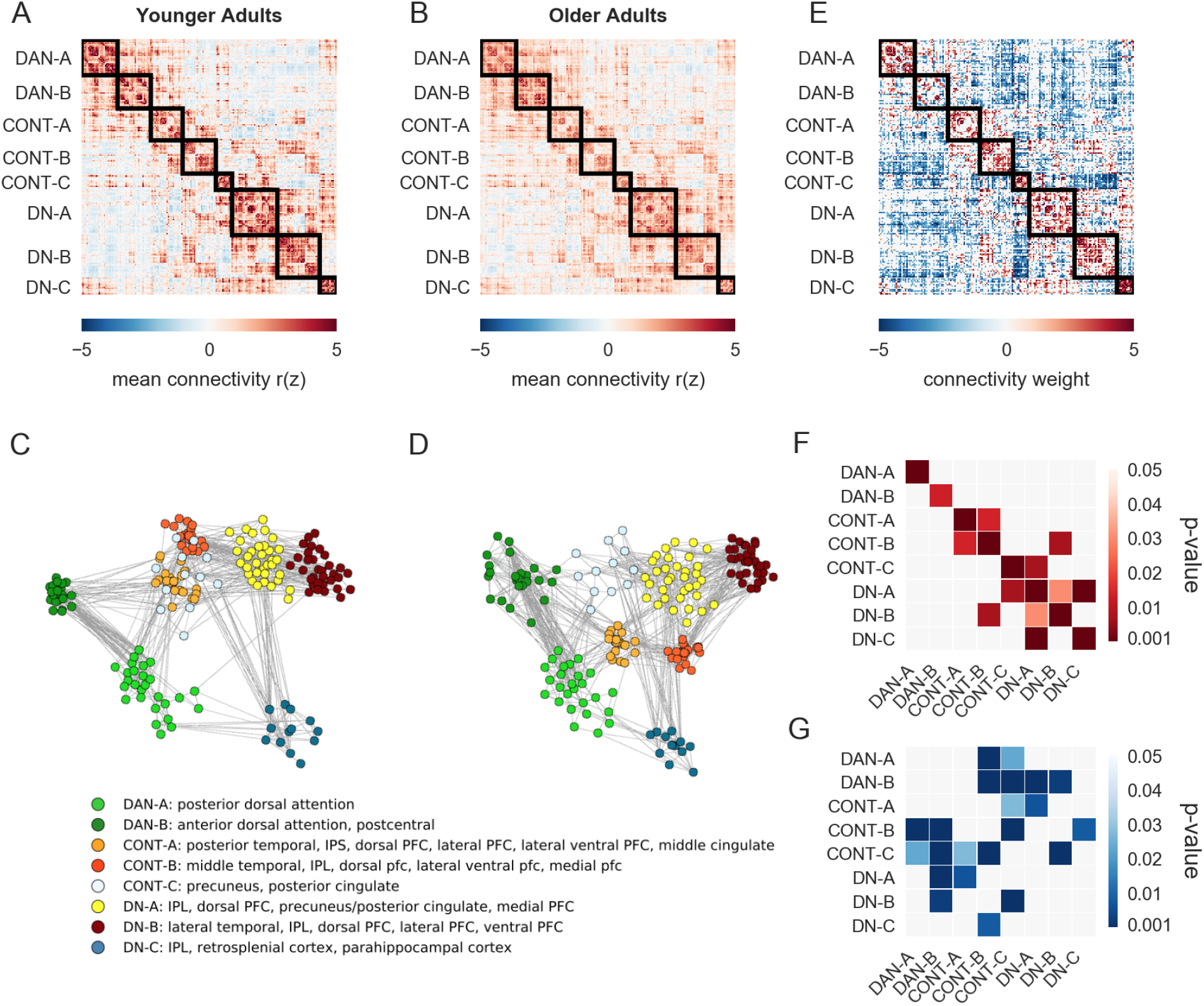
Functional connectivity of the default (DN), dorsal attention (DAN), and frontoparietal control (CONT) sub-networks following the Yeo 17-network solution. Mean group connectivity in (A) younger and (B) older adults. Spring-embedded plots (5% edge density) of the mean correlation matrices for (C) younger and (D) older adults. Nodes that are highly functionally correlated with one another are grouped closer together.(E) Differences in RSFC between younger and older adults among DAN, CONT, and DN. (F-G) Network contributions represent the summary of positive and negative edge weights within and between networks in younger (F) and older (G) adults. DAN = dorsal attention, FPC = frontoparietal control, DN = default.

Quantitative comparison with PLS of the inter-regional functional connectivity revealed a distinct pattern of age differences (permuted *p* < .001; Figure 5E). Younger adults (Figure 5F) showed more within-network connectivity. Between subnetwork connections were also seen in the young for CONT-A and CONT-B, and between DN-A to DN-B and DN-C. Between network connections in the young were also observed for CONT-B and DN-B. Older adults (Figure 5G) showed greater between-network connectivity of the dorsal attention network with frontoparietal control and default networks (DAN-A to CONT-B and CONT-C; DAN-B to CONT-B, CONT-C, DN-A, and DN-B), as well as greater frontoparietal control connectivity with the default network (CONT-A to DN-A; CONT-B to DN-C; CONT-C to DN-B). Older adults also showed greater connectivity among frontoparietal control subnetworks (CONT-A to CONT-C; CONT-B to CONT-C). A similar pattern of connectivity was observed with a 200 parcellation scheme (Supplemental Figure 9). Sub-network brain connectivity scores’ associations with cognition are reported in Supplemental Tables 2, 3, and 4, and Supplemental Figure 12.

### Connectomics Site Replication

To verify that our edge-level results are robust and replicable, and not confounded by potential overfitting of the PLS model, the full and sub-network PLS analyses were conducted only on the Ithaca sample. Brain connectivity scores were then computed from the Ithaca sample-derived weights and the Toronto sample individual-subject RSFC matrices, and compared between groups. Age group differences were replicated in the held out Toronto sample (*t*(61)= 6.42, *p* < .001, Cohen’s *d*= 1.63; *F*(1,57)= 21.13, *p* < .001, η_p_ = .27 with sex, education, and eWBV covariates included). In the sub-network analysis, age group differences were also replicated in the held out Toronto sample (*t*(61)= 7.01, *p* < .001, Cohen’s *d*= 1.79; *F*(1,58)= 24.16, *p* < .001, η_p_ =.29 with sex, education, and eWBV covariates included). These site replication analyses (Supplemental Figure 13) demonstrate that the PLS results are robust to potential issues of model overfitting and that the edge-level effects of functional brain aging observed in the Ithaca sample were also observed at the Toronto site.

### Cognition

Overall, predicted age-group differences in cognition were observed. Younger adults performed better on indices of episodic memory (*t*(281)= 17.51 *p* < .001; Cohen’s *d* = 2.11), executive function (*t*(281)= 12.67, *p* <.001; Cohen’s *d* = 1.52), and processing speed (*t*(281)= 15.03, *p* < .001; Cohen’s *d* = 1.81). Older adults had higher semantic memory index scores (*t*(281)= 9.18, *p* < .001; Cohen’s *d* = 1.10; see Table 1). Effects remained when testing for age group differences with ANCOVAs controlling for site, sex, education, and eWBV (Episodic: *F*(1,277)= 194.07, *p* < .001, η_p_^2^ = .41; Semantic: *F* (1,277)= 37.55, *p* < .001, η_p_^2^ = .12; Executive Function: *F*(1,277)= 132.70, *p* < .001, η_p_^2^ = .32; Processing speed: *F*(1,277)= 97.21, *p* < .001, η_p_^2^ = .26).

Associations between cognition and BOLD signal dimensionality, manifold eccentricity, and brain connectivity scores from the whole brain and sub-network analyses were examined (See Supplemental Tables 2, 3, and 4, and Supplemental Figures 10, 12 and 14). While several significant brain-behavior associations were observed, all of these fell below statistical significance thresholds after site was added as a covariate in the models.

## Discussion

Brain aging is marked by dedifferentiation in patterns of brain activity and functional connectivity. Here we adopted a comprehensive, multi-method approach to examine patterns of intrinsic network dedifferentiation across multiple spatial scales. Specifically, we applied novel methods to identify global, macroscale gradient, and edge-level differences in RSFC between younger and older adults. BOLD dimensionality, the number of BOLD (i.e., non-noise) components in the ME-fMRI signal, was lower for older adults, signaling a global shift towards dedifferentiated brain networks in older age. In contrast, the organization of macroscale connectivity gradients was largely preserved with age. However regional and global differences in connectivity gradients did emerge. Edge-level, multivariate analyses with PLS also revealed regional and network-specific patterns of dedifferentiation in older adulthood. Across the full cortical connectome, visual and somatomotor regions were more functionally integrated with other large-scale networks for older versus younger adults. In a targeted, sub-network analysis including default, dorsal attention, frontoparietal control networks, older adults showed greater default-executive coupling and reduced anticorrelation between default and dorsal attention networks. By examining age differences in the functional connectome across multiple spatial scales, we revealed that the intrinsic network architecture of the aging brain is marked by both global as well as topographically-discrete, network-specific patterns of functional dedifferentiation. The findings provide evidence for both global and network-specific patterns of dedifferentiation, laying the foundation for future studies examining alterations in RSFC as putative sensitive and specific markers of neurocognitive aging.

### BOLD signal dimensionality and global network dedifferentiation

Dimensionality in the BOLD signal was significantly lower for older versus younger adults, reflecting a generalized pattern of network dedifferentiation continuing into later life. This finding builds upon an earlier report of cross-sectional dimensionality reductions from adolescence to early and middle adulthood (Kundu et al., 2018; Figure 2). Reductions in dimensionality in early adult development, largely attributable to functional integration among prefrontal and other transmodal cortices, reflects the transition from local connectivity to longer-range connections and the formation of spatially distributed yet intrinsically coherent brain networks (Kundu et al., 2018). The shift in functional brain organization parallels cognitive development over this period, which is marked by the emergence of more integrative and complex cognitive functions (Zelazo and Carlson, 2012), and is also evident within the structural connectome (Park et al., 2020).

Declines in the dimensionality of the BOLD signal, which begin in adolescence, continue unabated throughout adulthood and into later life. In younger adults, lower dimensionality reflects greater functional integration and the emergence of large-scale brain networks (Kundu et al., 2018). However, our observation of continued reductions in BOLD signal dimensionality into older adulthood suggests that network integration may reach an inflection point in middle age (Zonneveld et al., 2019). After this point, continued reductions in dimensionality may no longer be driven by network integration, but rather by global network disintegration, and associated loss of coherent network components in the BOLD signal. Critically, our findings using this novel metric of BOLD signal dimensionality are consistent with earlier reports of age-related decreases in network modularity (Geerligs et al., 2015) and network segregation (Chan et al., 2014). Indeed these measures are reliably and positively correlated with dimensionality in our sample (see Supplemental Table S1). However, unlike these two graph analytic measures of network organization, BOLD signal dimensionality is agnostic with respect to the selection of cortical parcellation schemes, network definitions or specific network metrics. As such, we suggest that dimensionality may serve as a useful, data-driven marker of functional brain health in later life. An important next step in this regard will be to improve our mechanistic understanding of dimensionality reductions with age. Such global shifts may result from systemic structural, neurophysiological, metabolic or cerebrovascular changes known to occur with advancing age (e.g., Tsvetanov et al., 2020).

Finally, as a novel metric applied to a healthy aging sample, we acknowledge that there are important future directions to more fully interrogate the validity and applicability of BOLD signal dimensionality as an informative marker of functional brain aging. Additional work is necessary to conduct a validation of this metric, following the roadmap outlined by the original validation studies in younger and middle-aged adults (Kundu et al., 2013, 2017, 2018). As a physical property of T2* signal decay, the TE-dependence of BOLD signal (the fundamental component of ME-ICA BOLD signal denoising) should be largely robust to age differences. Directly testing this assumption will be an important direction for future research. Taken together, the TE-dependence of the BOLD signal as well as the validation studies conducted in healthy younger samples, give us confidence in BOLD signal dimensionality as a reliable, informative marker of brain aging.

### Gradients and macroscale connectomics

Reductions in BOLD signal dimensionality into older age suggest a global shift towards a dedifferentiated network architecture. We investigated whether this global shift may comprise more precise topographical patterns, reflected as greater similarity in connectivity profiles among brain regions. We tested this hypothesis by examining macroscale connectivity gradients in younger and older adults. While this is the first report of gradient analyses using ME-fMRI and individualized parcellation methods, our findings largely recapitulate connectivity gradients observed in young adults (Margulies et al., 2016). Transitions in functional connectivity patterns were observed from sensory/motor to transmodal association cortices (principal gradient) and from visual to somatomotor cortices (second gradient). This gradient architecture was similar for young and old, suggesting the macroscale organization of the gradients is generally preserved with age, as has been observed previously (Bethlehem et al., 2020). However, specific age-related regional differences did emerge in both gradient maps.

Age-related differences across both gradients included regions within visual, somatomotor and attentional networks. Differences within these clusters suggest a reduction in differentiation with respect to their corresponding gradient anchor (unimodal or transmodal in the case of the principal gradient, somatomotor or visual in the case of the second gradient). Specifically, both the superior parietal lobule, a node of the dorsal attention network implicated in externally-directed attention and visuomotor control processes, and somatomotor regions, showed greater similarity in connectivity profiles to transmodal regions. This is consistent with earlier reports, and patterns observed in the present edge-level analysis, of reduced anticorrelation between the dorsal attention and default networks in later life (Spreng et al., 2016).

Along the principal gradient, visual regions were more prominently anchored along the unimodal axis in older adults. This finding is consistent with the spring embedding plots of RSFC (Figure 4C-D), with more isolated visual regions among the top five percent of RSFC. This suggests that the principal gradient in older adults is likely more driven by the differentiation of heteromodal from visual systems, while in the younger adults there is a more marked differentiation of heteromodal from somatosensory/motor systems along this axis. This finding stands in contrast to the edge-level results (discussed below) which show greater age-related integration of visual and somatomotor cortices with heteromodal association regions. Importantly, gradients do not index functional connectivity strengths per se, but rather low dimensional patterns of RSFC between specific regions and the rest of cortex. Thus, the shift in connectivity profiles towards visual regions does not address functional integration of these regions, but their weight along the gradient axis. We suggest that thresholding the connectivity matrices before gradient mapping may have contributed to the age-related shift of visual cortices towards the unimodal anchor of the first gradient. While speculative, we suspect that matrix thresholding may have driven the suprathreshold connections towards an over representation of stronger local versus weaker long-range functional connections. This would be consistent with relative age-related reductions in volume and integrity of long-association fiber pathways versus local connections in primary sensory regions (Kochunov et al., 2012; Raz et al., 2005). These age-related structural differences would, in turn, render functional connectivity profiles towards more localized patterns, yielding an age-related shift in the gradient map towards the unimodal anchor. Nevertheless, these findings highlight the importance and potential impact of thresholding decisions, a point we return to in our discussion of edge-level connectivity below.

The gradient manifold, a scatter plot of the first two connectivity embedding gradients, represents organization across the functional hierarchies (Figure 3C). Manifold eccentricity depicts the Euclidean distance of regions from the center of the manifold, and displays the veridicality of the first against the second gradient. In older adults we observed greater diffusion of the vertices in the manifold with higher levels of eccentricity. Not limited to visual or somatosensory regions, the entire manifold is more diffusely organized, suggesting global dedifferentiation. This observation is consistent with previously observed increases in manifold dispersion across the lifespan (Bethlehem et al., 2020). This metric of manifold eccentricity was negatively correlated with the global measure of BOLD dimensionality, within and across groups. Lower BOLD dimensionality, as observed in older adults, was related to higher manifold eccentricity, providing cross-method convergence for global dedifferentiation.

It is important to note that we applied diffusion map embedding, a non-linear dimensionality manifold learning technique from the family of graph Laplacians (Coifman et al., 2005). This approach is among the most widely implemented in the literature (e.g. Bethlehem et al., 2020; Hong et al., 2019; Margulies et al., 2016; Murphy et al., 2019; Vos de Wael et al., 2020). However, given the novelty of gradient mapping in older adult populations, a direction for future research will be to critically evaluate the full range of approaches as well as algorithm parameters (Hong et al., 2020), including incorporation of repulsion properties (Böhm et al., 2021) in the gradient analysis to yield greater clarity regarding the segregation of discrete networks and differences with age.

### Edge-level connectomics

To more precisely investigate edge-level connectivity patterns, we adopted a multivariate analytical approach. As PLS uses singular value decomposition to test age differences across all edges in a single analytical step, we report RSFC differences across the full functional connectome, eliminating the need to apply functional connectivity strength or density thresholds. Visual inspection of the full connectomes for younger and older adults (Figure 4, Panels A-D) revealed a global pattern of network dedifferentiation for older adults, consistent with our dimensionality findings and previous reports (Betzel et al., 2014; Chan et al., 2014; Geerligs et al., 2015; Malagurski et al., 2020; Stumme et al., 2020). These qualitative differences were statistically validated in the group analysis (Figure 4, Panel E) and aggregate network matrices (Figure 4, Panels F-G). As predicted, younger adults showed a robust pattern of within-network connectivity, as well as connectivity between transmodal networks (Bullmore and Sporns, 2009; Gratton et al., 2012).

Despite preserved macroscale gradients, edge-level analyses revealed striking age differences in network-specific connectivity patterns. First, within-network connectivity was lower for older adults across the seven canonical networks investigated here. Reduced within-network functional connectivity is a hallmark of normative aging (Damoiseaux, 2017 for a review). We speculate that degraded within-network coherence is likely a key determinant of reduced BOLD signal dimensionality, and global network dedifferentiation, in older adulthood. In addition to lower within-network coherence, edge-level analyses also revealed three distinct, network-specific dedifferentiation patterns. The most striking of these revealed greater integration of visual and somatomotor regions with all other networks for older adults (Figure 4, Panel G). Functional integration of visual and somatosensory regions has been observed previously. Chan and colleagues (2014) reported reduced segregation of visual cortices from other brain networks, although this was not explicitly quantified in their analyses. Similarly, age-related increases in node participation, a graph analytic marker of functional integration, were limited to visual and somatosensory networks in a large study of age differences in RSFC (Geerligs et al., 2015). Further, Stumme and colleagues (2020) reported that age differences in RSFC were most prominent in visual and somatosensory cortices. While previous studies reported patterns of sensorimotor integration with age, these have not gained prominence as a central feature of functional brain aging. As discussed above with regards to the gradient analysis results, statistical thresholding of the gradient matrices might significantly impact these findings. Threshold-based approaches highlight age-related differences among the most robust connections, often associated with heteromodal cortices, potentially obscuring less robust age differences in other networks. This is particularly evident in our findings, where somatomotor and visual networks show small age differences relative to those observed for association networks in the thresholded, spring-embedded plots (Figure 4, panels C-D). In contrast, analysis of the unthresholded matrices revealed integration of sensorimotor networks to be among the most striking features of the aging connectome (Figure 4, panels E-G).

Our findings of greater visual network integration parallel task-based studies identifying greater top-down modulation of visual association cortices by transmodal regions as a central feature of functional brain aging. Greater activation of transmodal cortices, in the context of age-related declines in the fidelity of sensory signaling, has been interpreted as increased demand for top-down modulation of early sensory processing (Clapp et al., 2011; Li and Rieckmann, 2014; Payer et al., 2006; Spreng and Turner, 2019b). Indeed, sensory declines and motor slowing account for much of the individual variability in cognitive functioning among older adults (Baltes and Lindenberger, 1997; Salthouse, 1996). This suggests that greater modulation of these primary sensorimotor regions (and visual attention and visuomotor control regions of the superior parietal lobule, see ‘Gradient analyses’ above) may be necessary to sustain complex thought and action in later life. While beyond the scope of the current study, we speculate that such task-driven demands for greater cross-talk between transmodal and sensorimotor cortices may, in turn, shape the intrinsic functional architecture of these networks in older adulthood (Stevens and Spreng, 2014).

We also conducted a targeted analysis of edge-level age differences in the default, dorsal attention, and frontoparietal control networks. Previous work has demonstrated that these networks interact during goal-directed cognitive tasks (Spreng et al., 2010; Dixon et al., 2018; Murphy et al., 2020), show similar connectivity profiles during both task and rest (Spreng et al., 2013) and undergo significant changes into older adulthood (Grady et al., 2016; Sullivan et al., 2019; Ng, et al., 2016). For this *a priori* analysis, we adopted the sub-network topography for the three networks derived from the 17-network solution (Yeo et al., 2011). This enabled us to investigate age-related changes with greater precision. Importantly, as we observed for the full connectome analysis, the thresholded spring-embedded plots (Figure 5, panels C-D) failed to reveal the robust age-differences in connectivity among default, dorsal attention and frontoparietal control network regions that emerged from the edge-level analyses (Figure 5, panels E-G). While the predicted pattern of reduced within-network connectivity was recapitulated across the sub-networks, we observed two additional network-specific dedifferentiation patterns. As predicted, there was greater age-related coupling of default and frontal brain regions, a pattern we have described as the Default to Executive Coupling Hypothesis of Aging (DECHA; Turner and Spreng, 2015; Spreng and Turner, 2019a). This pattern did not emerge in the seven network analysis (Figure 4). However, when applied to the edge-level sub-network matrices (Figure 5, panels E-G), a clear DECHA pattern emerged for CONT-A to DN-A, CONT-B to DN-C, and CONT-C to DN-B sub-networks (Figure 5, panel G). While we did not identify reliable associations with cognition here, we have posited that this dedifferentiation pattern may reflect the shifting architecture of cognition in later life (Turner and Spreng, 2015; Spreng et al., 2018) with both adaptive and maladaptive consequences for cognitive aging (Spreng and Turner, 2019a).

A second dedifferentiation pattern emerged in this sub-network analysis. Older adults showed greater connectivity between the dorsal attention and the two other association networks. This pattern was particularly pronounced for the DAN-B sub-network which includes the superior parietal lobule. Previous reports have shown reduced anticorrelation between dorsal attention and default networks (Keller et al., 2015; Spreng et al., 2016) in older adulthood. These edge-level findings also converge with our gradient analyses where the superior parietal lobule, a node of DAN-B, showed an age difference in connectivity gradient, with a functional connectivity profile more similar to that of other transmodal regions. The DAN-B sub-network encompasses regions of the putative frontal eye fields and precentral gyrus implicated in top-down, or goal-directed, attentional control. This is again consistent with a neuromodulatory account of neurocognitive aging, wherein greater allocation of attentional resources may be engaged to sharpen perceptual representations in later life (Li et al., 2006; Li and Rieckmann, 2014). Future research will be necessary to directly test these hypotheses, linking network-specific patterns of dedifferentiation to domain specific cognitive changes.

### Cognitive Function

Our findings suggest that both global and network-specific dedifferentiation are core features of the functional aging connectome. In a final series of analyses we investigated whether these network changes were associated with cognitive functioning. We observed significant behavioral correlations with BOLD signal dimensionality and edge-level connectivity. Intriguingly however, all observed associations fell below statistical significance thresholds when site was included as a covariate in the statistical models. This was the case even though both brain and behavioral age effects replicated across both sites (see Supplemental Figure 13 and Supplemental Table 5). As a result we do not interpret the brain and behavior associations further here and report all uncorrected and partial correlations in Supplemental Tables 2, 3 and 4 (see site-specific scatter plots Supplemental Figure 14). While we took extraordinary care to match data collection protocols and core demographics, study site encompasses many additional moderating factors that may have influenced brain and behavioral associations across the two sites (e.g., socioeconomic status, see Chan et al., 2018). While increases in statistical power enabled by multi-site investigations permit greater sensitivity to detect brain-behavior associations, it also comes at the potential cost of structured noise related to population differences. Understanding these differences will also be an important direction for future research.

## Conclusion

We employed a multi-method data acquisition and analysis protocol to study functional brain aging across multiple spatial scales, with a specific emphasis on age-related patterns of intrinsic network dedifferentiation. Reduced BOLD signal dimensionality suggested a global, age-related shift towards dedifferentiated network organization in older versus younger adults. Limitations of a cross-sectional study design restrict interpretations with respect to lifespan shifts in brain function. However, we speculate that network integration across the adult lifespan may include an inflection point in middle adulthood, beyond which network integration in early adulthood shifts to a pattern of network dedifferentiation, and the dissolution of a segmented and modular network architecture.

The methodological and analytical approach adopted here were selected to, at least in part, overcome several of the most enduring and pervasive challenges in lifespan network neuroscience. These include age-related variability in noise profiles within the BOLD signal, as well as distortions introduced by group-wise spatial alignment to standardized templates. Of course the methods implemented here cannot address the totality of confounds that complicate RSFC analyses. Among the most critical of these, and an important direction for future research, is resolving, or at least accurately modeling, age differences in neurovascular coupling. Altered neurovascular coupling with age can introduce spurious RSFC differences that are difficult to detect with standard imaging protocols (Tsvetanov et al., 2020). While ME-ICA methods, which separate neural from non-neural sources in the BOLD signal, are a significant advance, implementation of multimodal methods such as simultaneous arterial spin labeling and echo-planar imaging may be necessary to resolve this issue (Tsvetanov et al., 2020). Additionally, residual motion-related noise was still observed in the BOLD signal, which could be attributable to respiration (e.g. Power et al., 2018; Lynch et al., 2020). While this noise did not confound our age effects, its persistence requires additional consideration and points to the need for further advances to improve signal-to-noise with ME-fMRI data. Despite these limitations, we suggest that the multifaceted approach adopted here offers a comprehensive account of age differences in the functional network architecture of the brain, including both novel and previously observed patterns of network dedifferentiation and integration. Taken together, these findings add further clarity and precision to current understanding of how functional networks are formed, shaped, and shifted into older adulthood.

## Acknowledgments

This work was supported in part by NIH Grant 1S10RR025145 and by a Canadian Institute of Health Research grant to R.N.S..

## Competing Interests

The authors declare no competing interests.

## Supplemental Material

**Supplemental Figure 1:**
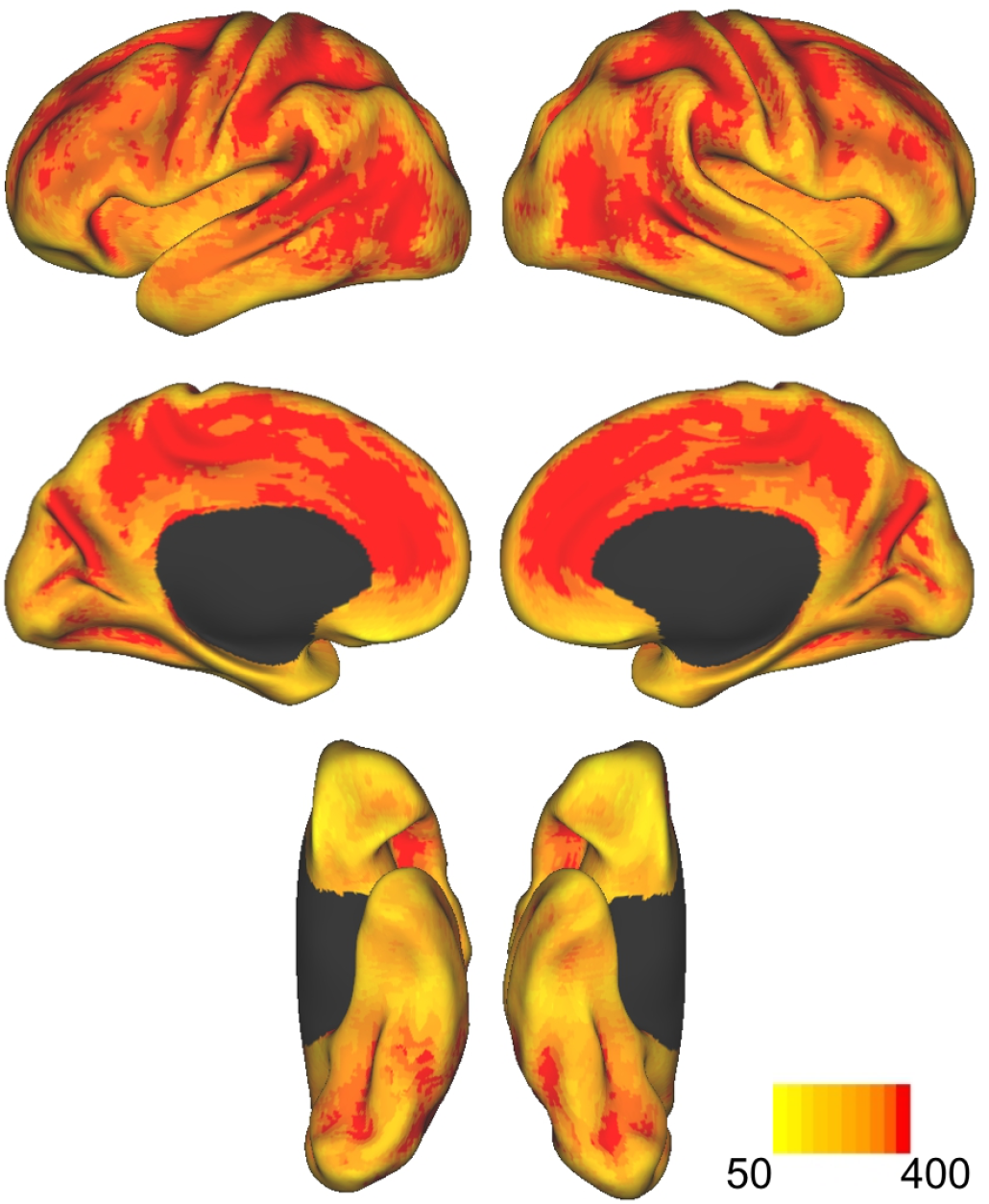
Temporal Signal to Noise. A temporal signal to noise ratio (tSNR) map was created for both runs of each participant’s denoised data and averaged. The average group map (N=301) was projected onto an inflated surface, separated by left and right hemispheres. Maps were thresholded to demonstrate that the low end of tSNR values were well above 50. A maximum tSNR of 400 is shown here for visualization purposes.

### Impact of Motion on RSFC

ME-ICA has been shown to effectively remove distant dependent motion effects in RSFC data (Power et al., 2018). To rule out the possibility that these residual motion effects confounded our main results, we conducted a behavior PLS analysis with raw framewise displacement (FD). This analysis yielded a significant pattern (59.21% covariance explained, permuted *p* = .004; Supplementary Figure 2A) representing the main effect of correlating functional connectivity with FD across younger and older adults (younger adults r = .70; older adults r = .65; Supplementary Figure 2B). While a motion-related connectivity pattern emerged (explaining 59.2% of the cross-block covariance), no significant group differences or age group interactions emerged. Network contribution plots expressing the mean positive and negative weights within and between networks are depicted in Supplementary Figure 2C. Higher FD was associated with greater within-VIS connectivity and LIM, FPC, and DN connectivity (warmer colors), as well as lower within-network connectivity of SOM, DAN, VAN, LIM, and DN (cooler colors).

As described in Methods, covariance in PLS can be specified as percent cross-block covariance, where the cross-block covariance is between the brain data and predictor variables. The sum of the percent cross-block covariance must sum to 100% over all latent variables (LVs). As such, the value of covariance explained for a given LV cannot be interpreted in isolation but must be weighed against the other LVs. This is demonstrated in the case of a single predictor variable, where the resultant LV will explain 100% of the cross-block covariance.

These findings suggest that ME-ICA patterns of RSFC are still impacted by motion. BOLD signal post-MEICA has been related to residual respiratory effects (Power et al., 2018), but to our knowledge there is no evidence to suggest that this residual noise would confound group comparisons. Rather, residual motion may reduce signal-to-noise in a similar manner across groups. Indeed, we find that motion effects are not confounded by group membership in our sample. A second LV accounting for 40.79% (p = .38) of the covariance dissociated age groups in their relationship to motion. Relative to the first LV, group specific motion effects on connectivity account for less covariance in the data. This LV, however, was not significant . If this LV was significant, then there would be evidence that motion differentially impacted connectivity by group, limiting the interpretation of the data. Because this interaction was not significant, our primary results, all of which compare groups, are not confounded by motion.

We empirically confirmed that the pattern of connectivity covarying with motion is not consistent with the age differences in RSFC reported in the main text. We determined this by assessing the correspondence between brain connectivity scores from the whole-brain age contrast (Figure 4) and from the motion-associated effects (Supplementary Figure 2A-C). Relationships are plotted in Supplementary Figure 2D-F. Partial correlations controlling for sex, education, eWBV, and site were as follows: Across the full sample, *pr* = -.037, p = .534; in younger adults: *pr* = -.049, p = .528; in older adults *pr* = .019, p = .833.

**Supplemental Figure 2:**
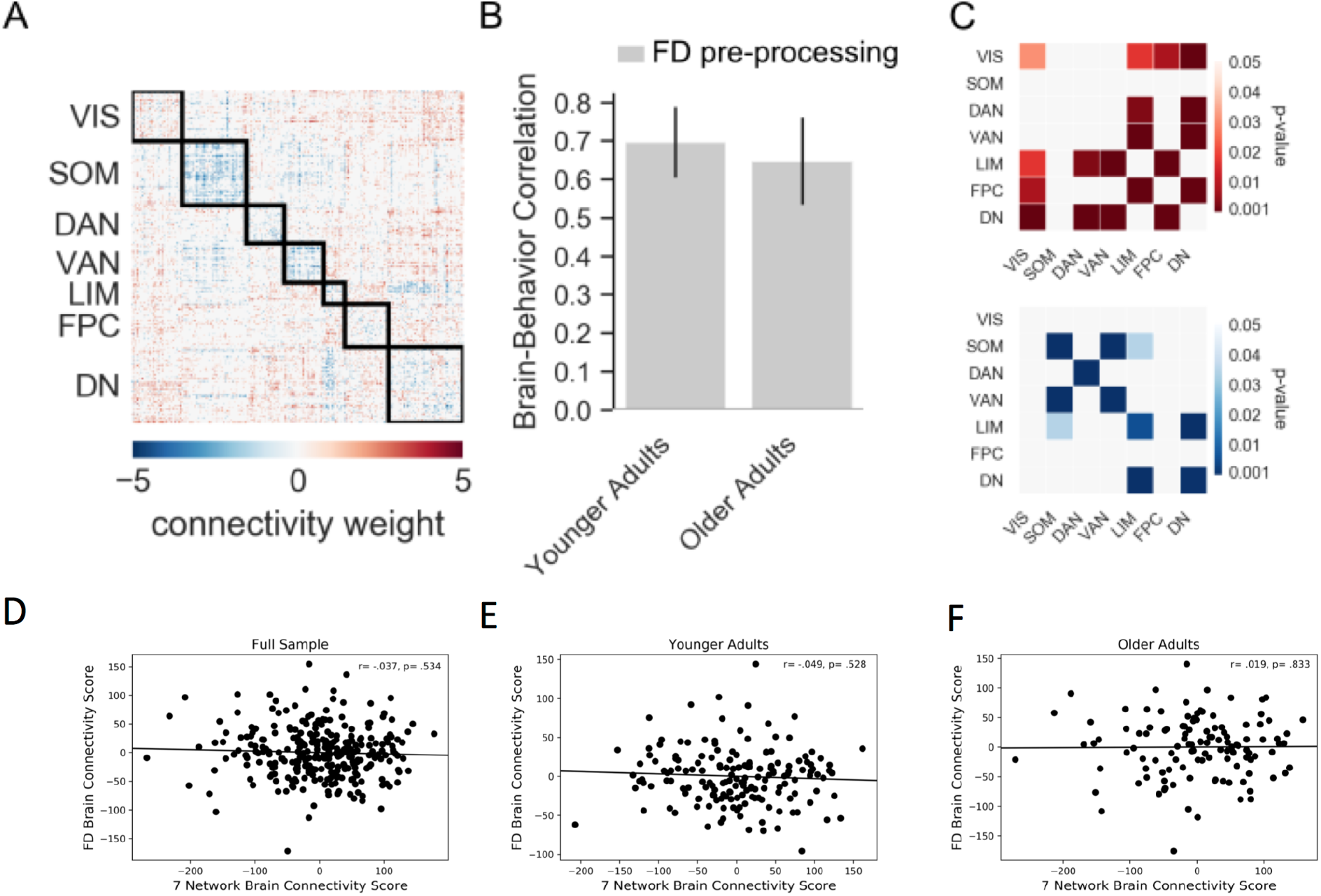
Behavioral PLS examining the relationship between functional connectivity with motion in younger and older adults. Framewise displacement (FD) was calculated on the middle echo prior to processing. (A) Functional connections that correlate with higher FD (warmer colors) and lower FD (cooler colors). (B) Correlations of FD with the functional connectivity pattern in younger and older adults. Error bars indicate the 95% confidence interval derived from the bootstrap estimation. (C) Network contributions of the mean positive and negative edge weights within and between networks. Scatterplots depicting age differences in brain connectivity scores on the x-axis and motion-related brain connectivity scores on the y-axis (D) across the full sample, (E) in younger adults, (F) in older adults. VIS = visual, SOM = somatomotor, DAN = dorsal attention, VAN = ventral attention, LIM = limbic, FPC = frontoparietal control, DN = default networks.

**Supplemental Figure 3:**
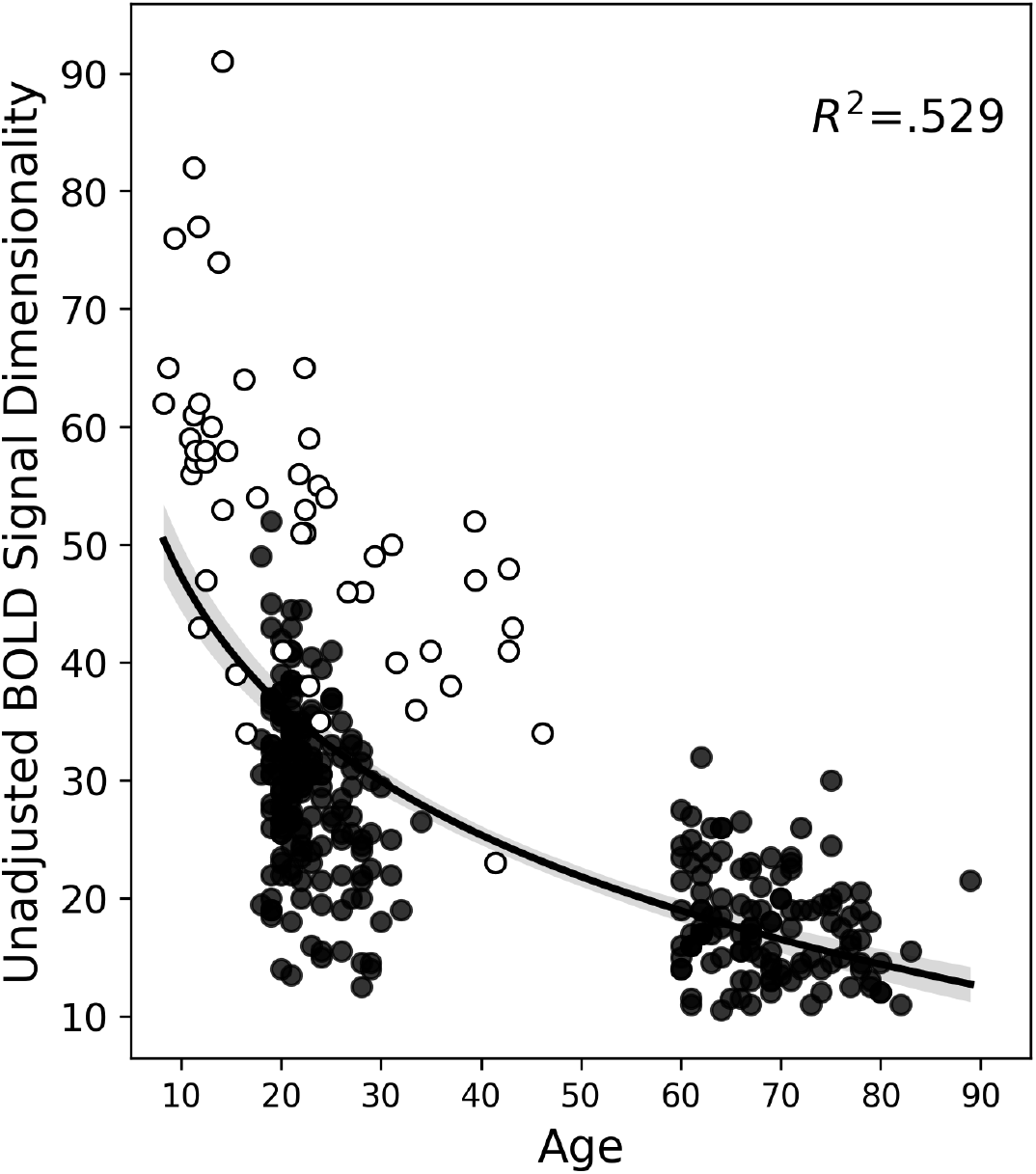
BOLD signal dimensionality. The scatter plot shows BOLD signal dimensionality by age with a power distribution and 95% confidence intervals overlaid. Points in white were contributed by Kundu and colleagues (2018). Here, BOLD dimensionality is not adjusted by the number of time points acquired. BOLD dimensionality was averaged across two runs for points in black.

### Characterizing BOLD Dimensionality

The present report leverages ME fMRI acquisition and processing to define a novel metric of BOLD dimensionality as a proxy for global functional integration. We further investigate the biological substrate of BOLD dimensionality by examining its most defining interregional connections and comparing it to similar summary statistics of global RSFC organization.

We first used behavior PLS to characterize the interregional connections associated with BOLD dimensionality. In brief, behavior PLS identifies functional connectivity patterns that optimally co-vary with a behavioral measure. Two significant patterns captured the association between functional connectivity and BOLD dimensionality in younger and older adults. The first pattern reflects a main effect of BOLD dimensionality across groups (69.69% covariance explained, permuted *p* < .001; younger adults r = .86; older adults r = .88; Supplementary Figure 4A). Network contribution plots summarize the significant interregional associations. Higher BOLD dimensionality was related to more within-network connectivity (Supplementary Figure 4A, warmer colors). Higher BOLD dimensionality was also related to connectivity among heteromodal association networks LIM, FPC, and DN . Dimensionality was negatively related to VIS connectivity across the connectome (Supplementary Figure 4A, cooler colors).

The second pattern revealed an age group difference in the association between functional connectivity and BOLD dimensionality (30.31% covariance explained, permuted *p* < .001; younger adults r = .62; older adults r = -.48). In younger adults, higher BOLD dimensionality was related to more connectivity within and between FPC and DN (Supplementary Figure 4B, warm colors). In contrast, higher BOLD dimensionality in older adults was related to more SOM connectivity with itself and with VIS, DAN, VAN, and LIM (Supplementary Figure 4B, cool colors).

**Supplemental Figure 4:**
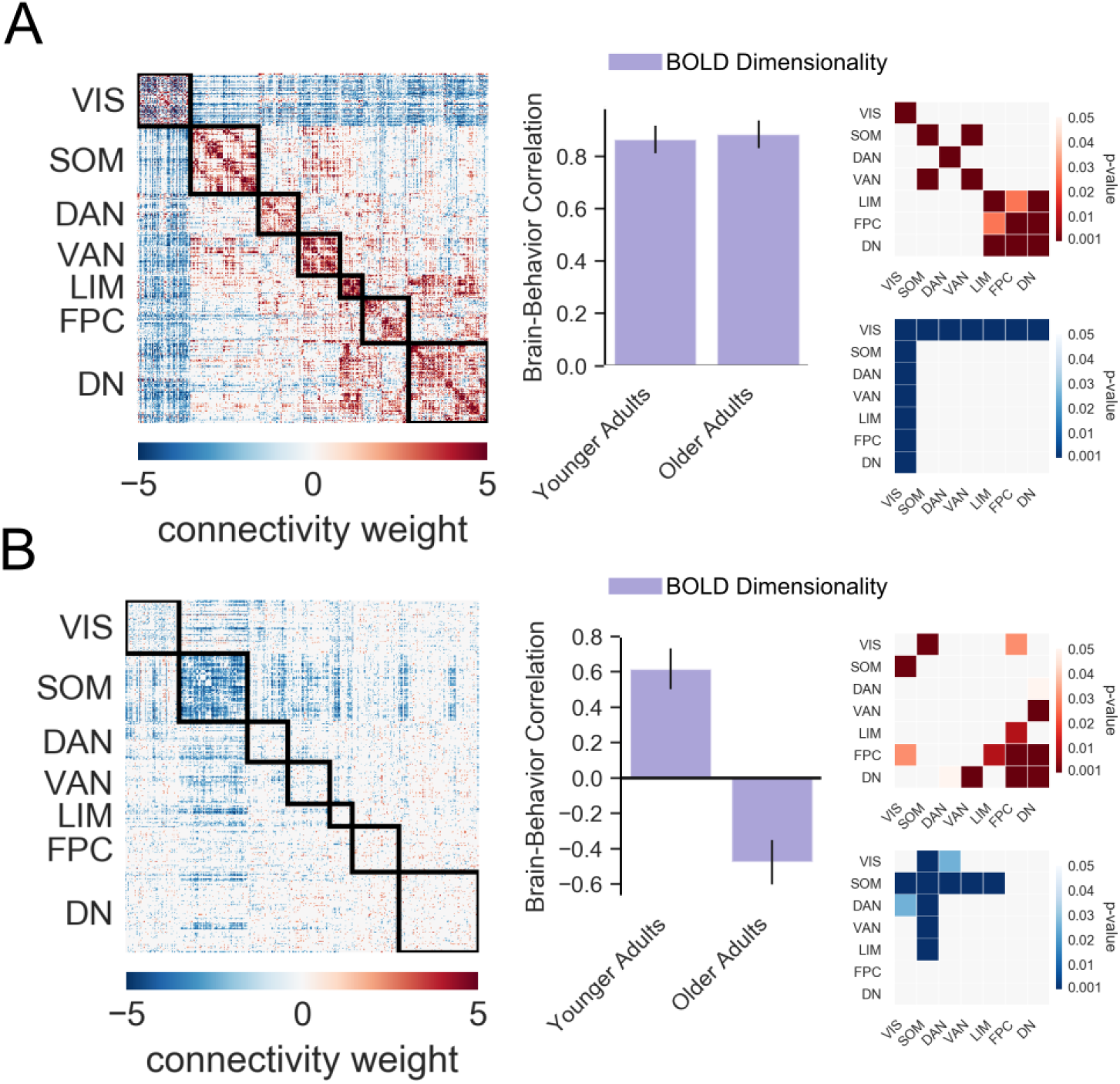
Behavioral PLS of the relationship between functional connectivity and BOLD dimensionality in younger and older adults. Two significant latent variables were identified. (A) The first latent variable expresses an effect of the number of BOLD components on functional connectivity. From left to right: functional connections that correlate with a higher (warmer colors) and lower (cooler colors) number of BOLD components; Bar plots showing the correlations of BOLD dimensionality with the functional connectivity pattern in younger and older adults; Network contributions of the mean positive (warm colors) and negative (cool colors) edge weights within and between networks. (B) The second latent variable expresses age-related differences in the association between functional connectivity and BOLD dimensionality. From left to right: Functional connections that correlate with a higher number of BOLD components in younger (warm colors) and older adults (cool colors); Bar plots showing the correlation between BOLD dimensionality and the functional connectivity pattern; Network contributions of the mean positive (warm colors) and negative (cool colors) edge weights within and between networks. Error bars for the brain-behavior barplots indicate 95% confidence intervals derived from the bootstrap estimation. VIS = visual, SOM = somatomotor, DAN = dorsal attention, VAN = ventral attention, LIM = limbic, FPC = frontoparietal control, DN = default.

We then compared BOLD dimensionality to a series of graph theoretic measures to determine whether it provides unique information about global integration. Participation coefficient, and modularity were computed as implemented in the Brain Connectivity Toolbox (Rubinov & Sporns, 2011). Segregation was also calculated (Chan et al., 2014). Measures were calculated on each individual’s z-scored functional connectivity matrix. Self-connections and negative weights were set to zero. Product-moment correlations were computed within each age group and on the full sample controlling for age, as shown in Supplemental Table 1 and Supplementary Figure 5. BOLD dimensionality was negatively associated with participation coefficient and positively associated with modularity and segregation in younger adults. In older adults, BOLD dimensionality was associated with segregation. In the full sample controlling for age, all graph metrics showed a significant relationship to BOLD dimensionality. The observed negative association between BOLD dimensionality and participation coefficient is consistent with a prior report of a negative association in a lifespan sample from children to middle-aged adults (Kundu et al., 2018). The positive association with modularity only in young adults reinforces that BOLD dimensionality highlights an inflection point in network organization across the lifespan: More dimensionality in young supports an established community structure; more dimensionality in older adults may support segregation as community structure devolves.

**Supplemental Table 1.** BOLD Dimensionality Associations with Graph Metrics.

| Metric | Younger Adults | Older Adults | Full Sample |
| --- | --- | --- | --- |
| Participation Coefficient | **-.491 (.000) [-.60, -.36]** | -.158 (.085) [-.33, .02] | **-.369 (.000) [-.47, -.26]** |
| Modularity | **.285 (.000) [.14, .42]** | .022 (.813) [-.16, .20] | **.129 (.030) [.01, .24]** |
| Segregation | **.689 (.000) [.60, .76]** | **.549 (.000) [.41, .66]** | **.536 (.000) [.45, .61]** |

*Note.* Metrics were derived from unthresholded correlation matrices containing positive connection weights. Partial correlations (*pr*) were calculated with sex, education, eWBV, and site as covariates. Age was included as an additional covariate in associations on the full sample. *r* values are presented with *p* values in parentheses and 95% confidence intervals in brackets. Bold denotes associations deemed significant at $p < .05$ .

**Supplemental Figure 5:**
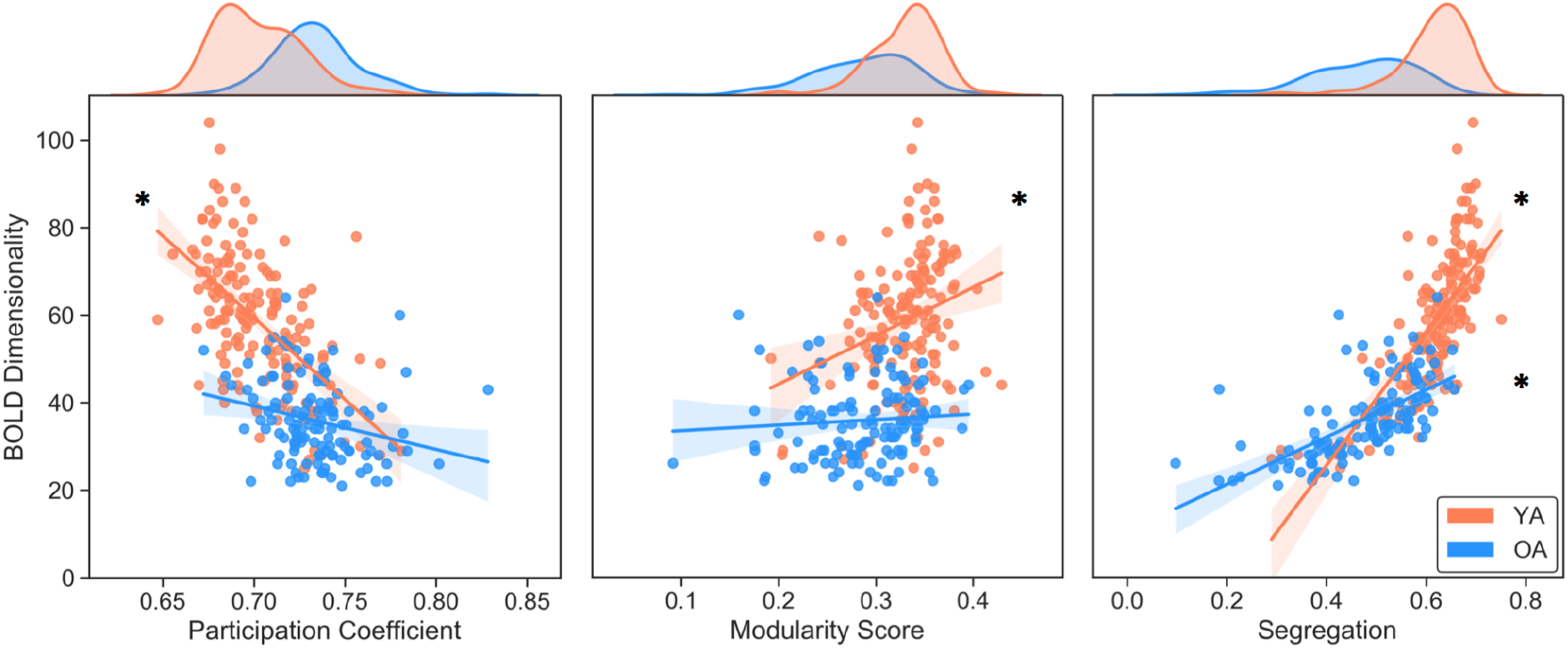
Scatterplots between BOLD dimensionality and graph theory measures. Distributions of graph measures are shown at the top of each plot. BOLD dimensionality distributions are shown in the rightmost plot. * indicates significant correlations. YA= younger adults; OA = older adults.

**Supplemental Figure 6:**
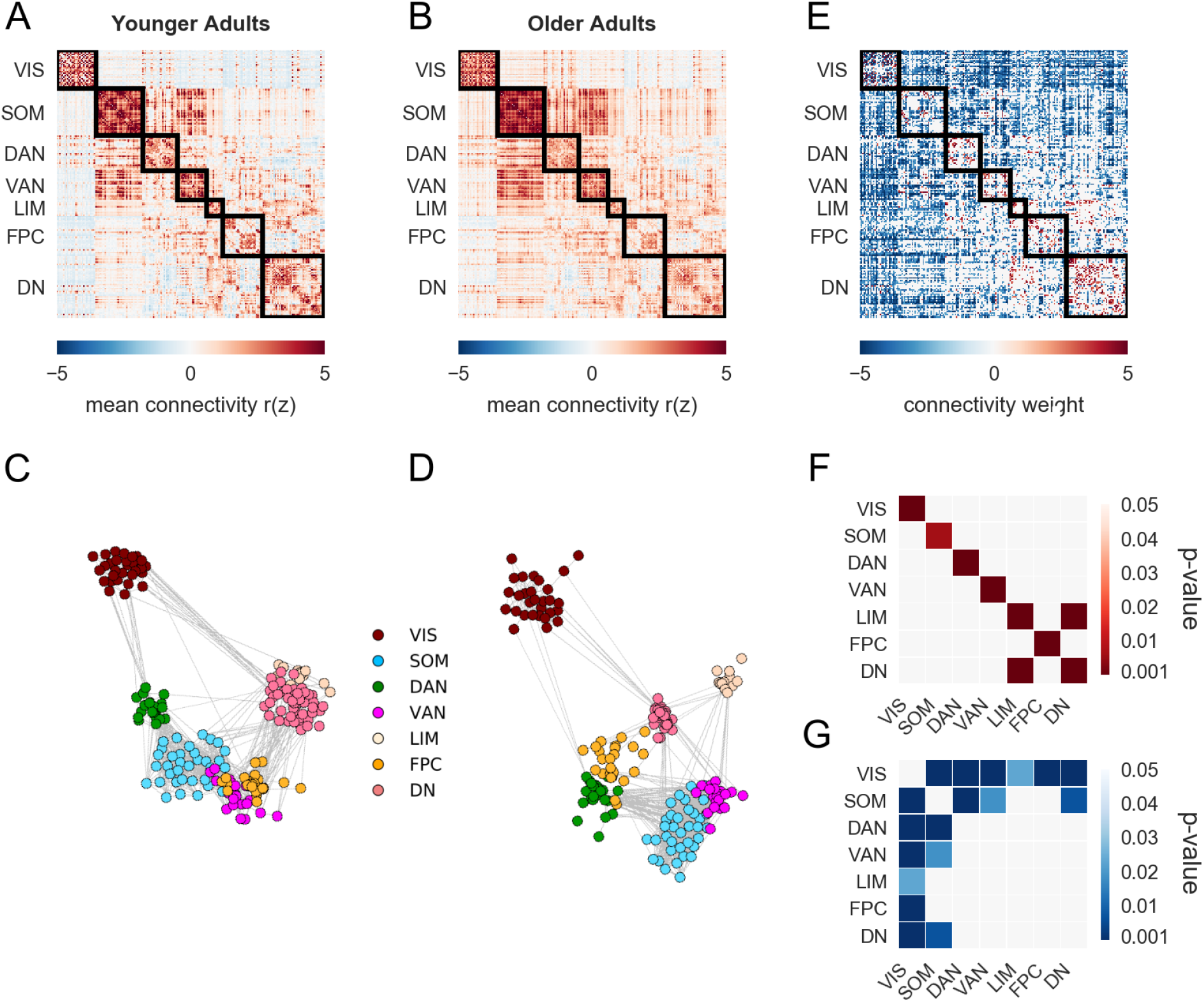
Functional connectomics in younger and older adults. Mean RSFC for the 200-parcellated MEFC data in (A) younger and (B) older adults. Spring-embedded plots with a 7-network solution (5% edge density) of the mean correlation matrices for (C) younger and (D) older adults. (E) Multivariate PLS analysis was used to identify age-related differences in RSFC between younger and older adults (p < .001). Red color indicates greater RSFC in younger adults, and blue color indicates greater RSFC in older adults. (F-G) Network contributions represent the summary of positive and negative edge weights within and between networks in younger (F) and older (G) adults. The mean positive and negative bootstrap ratios within and between networks are expressed as a *p* value for each z-score relative to a permuted null model. Higher values indicate greater connectivity than predicted by the null distribution. VIS = visual, SOM= somatomotor, DAN= dorsal attention, VAN = ventral attention, LIM = limbic, FPC= frontoparietal control, DN= default.

**Supplemental Figure 7:**
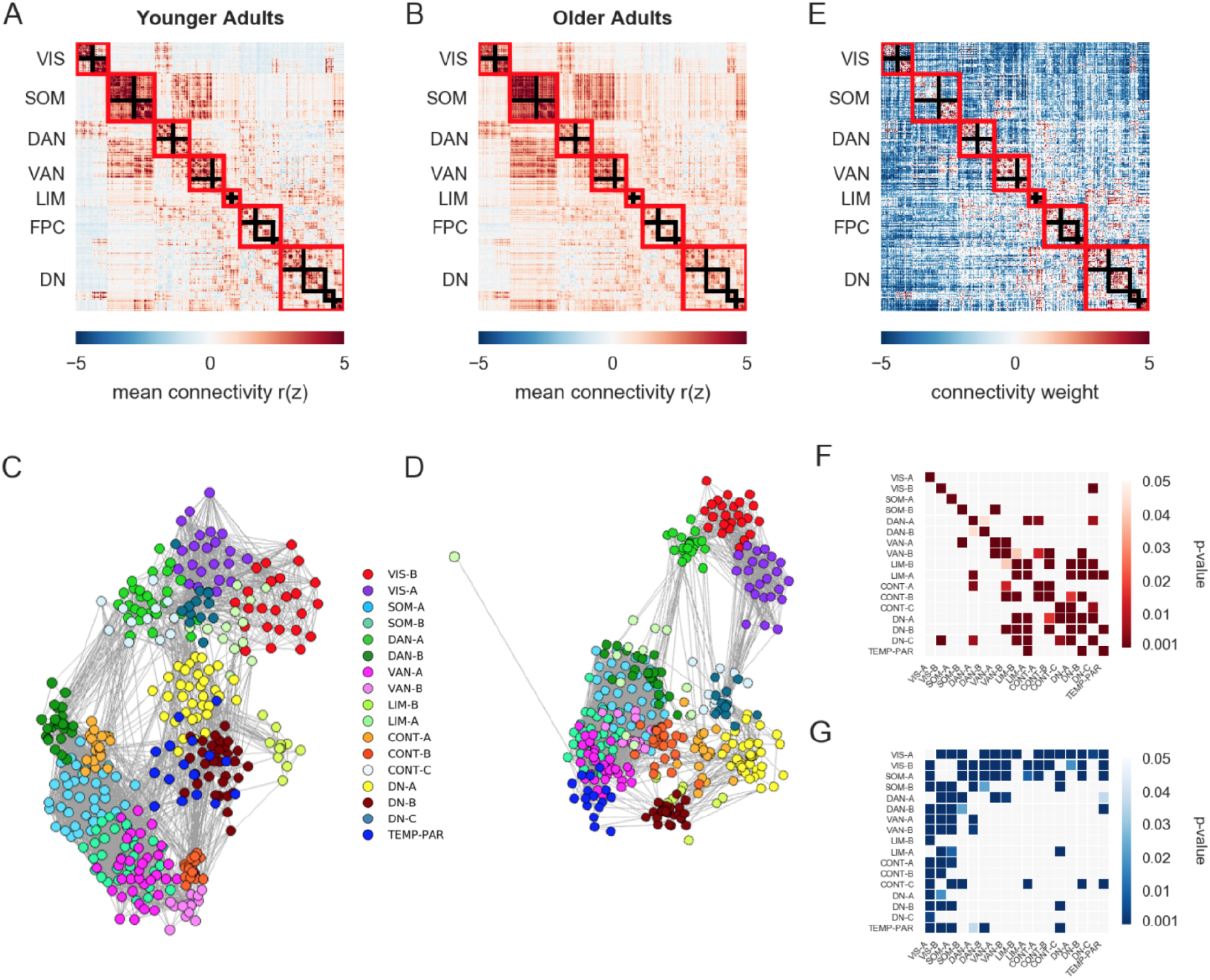
Functional connectomics in younger and older adults. Mean RSFC for the 400-parcellated MEFC data in (A) younger and (B) older adults. Spring-embedded plots with a 17-network solution (5% edge density) of the mean correlation matrices for (C) younger and (D) older adults. (E) Multivariate PLS analysis was used to identify age-related differences in RSFC between younger and older adults. Red color indicates greater RSFC in younger adults, and blue color indicates greater RSFC in older adults (p < .001). (F-G) Network contributions represent the summary of positive and negative edge weights within and between networks in younger (F) and older (G) adults. The mean positive and negative bootstrap ratios within and between networks are expressed as a *p* value for each z-score relative to a permuted null model. Higher values indicate greater connectivity than predicted by the null distribution. VIS = visual, SOM = somatomotor, DAN = dorsal attention, VAN = ventral attention, LIM = limbic, FPC = frontoparietal control, DN = default.

**Supplemental Figure 8:**
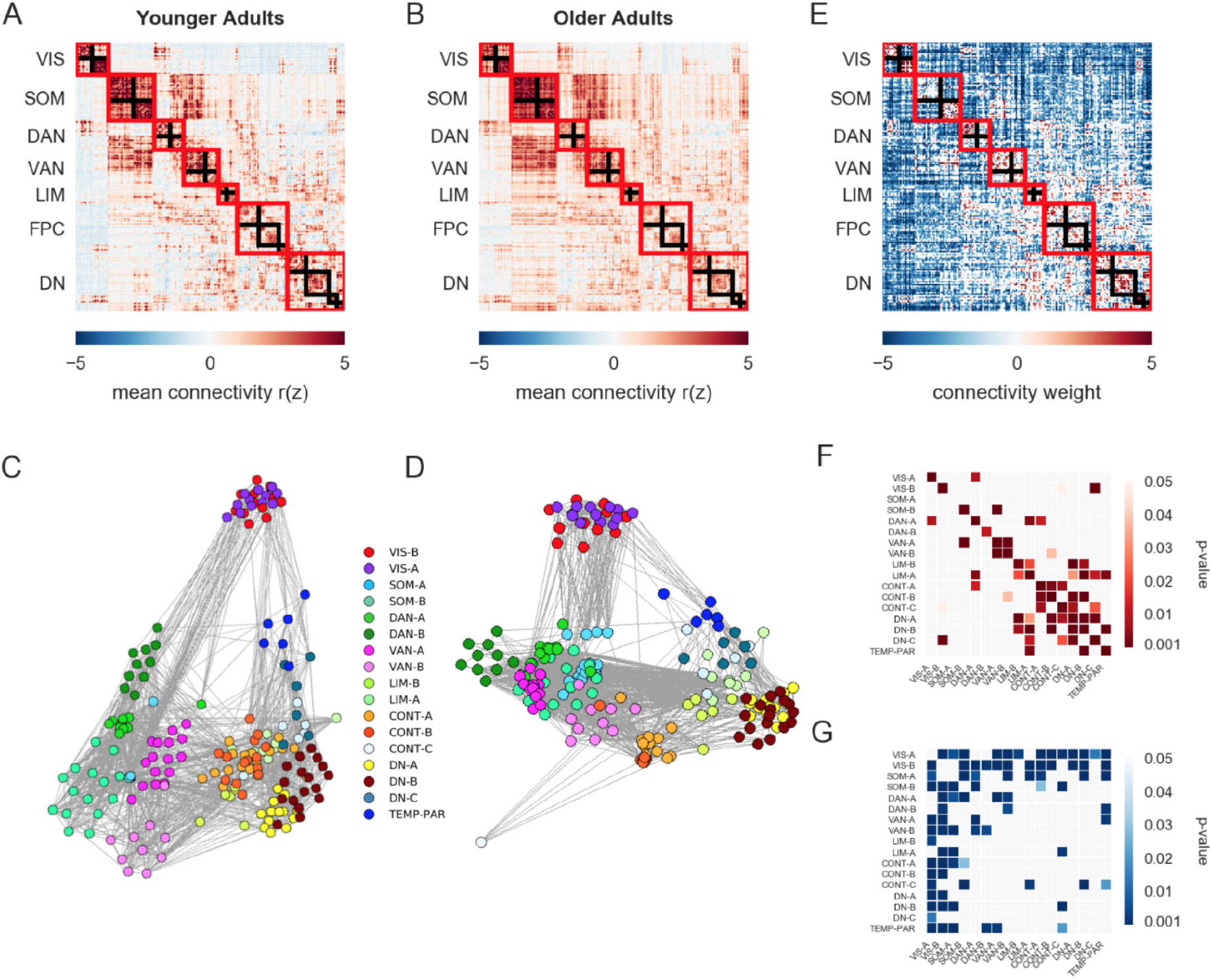
Functional connectomics in younger and older adults. Mean RSFC for the 200-parcellated MEFC data in (A) younger and (B) older adults. Spring-embedded plots with a 17-network solution (10% edge density) of the mean correlation matrices of (C) younger and (D) older adults. The threshold used for the spring-embedded plots was set higher due to the sparsity of the graph at thresholds lower than 10%. (E) Multivariate PLS analysis was used to identify age-related differences in RSFC between younger and older adults (p < .001). Red color indicates greater RSFC in younger adults, and blue color indicates greater RSFC in older adults. (F-G) Network contributions represent the summary of positive and negative edge weights within and between networks in younger (F) and older (G) adults. The mean positive and negative bootstrap ratios within and between networks are expressed as a *p* value for each z-score relative to a permuted null model. Higher values indicate greater connectivity than predicted by the null distribution. VIS = visual, SOM = somatomotor, DAN = dorsal attention, VAN = ventral attention, LIM = limbic, FPC = frontoparietal control, DN = default.

**Supplemental Figure 9:**
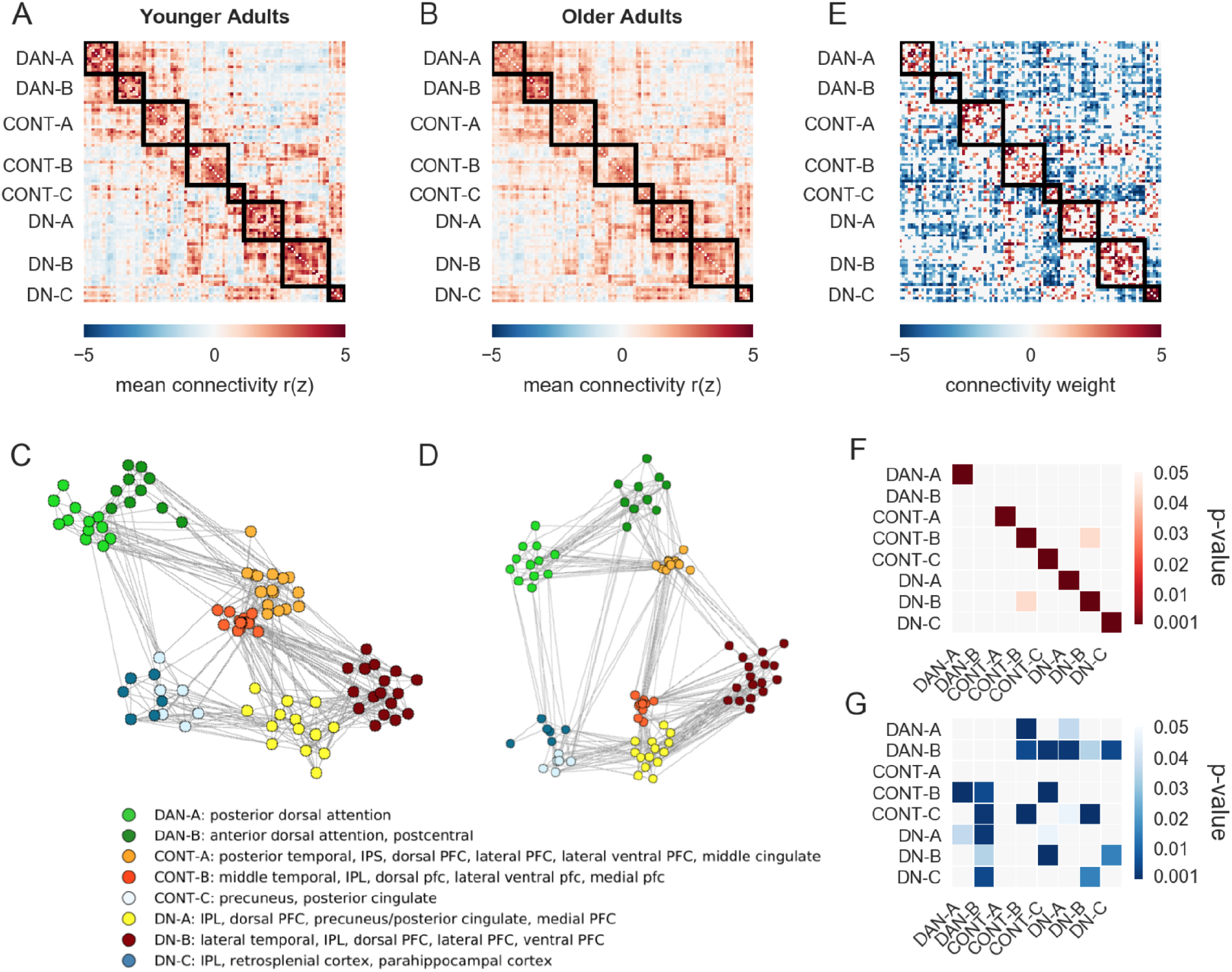
Functional connectivity of the default (DN), dorsal attention (DAN), and frontoparietal control (CONT) sub-networks following Yeo 17-network solution. Mean group connectivity for the 200-parcellated MEFC data in (A) younger and (B) older adults. Spring-embedded plots (10% edge density) of the mean correlation matrices for (C) younger and (D) older adults. A lower threshold was used for the spring-embedded plots due to the sparsity of the graph at higher thresholds. (E) Differences in RSFC between younger and older adults among DAN, CONT, and DN (*p* < .001). (F-G) Network contributions represent the summary of positive and negative edge weights within and between networks in younger (F) and older (G) adults. The mean positive and negative bootstrap ratios within and between networks are expressed as a *p* value for each z-score relative to a permuted null model. Higher values indicate greater connectivity than predicted by the null distribution. DAN = dorsal attention, FPC = frontoparietal control, DN = default.

**Supplemental Figure 10:**
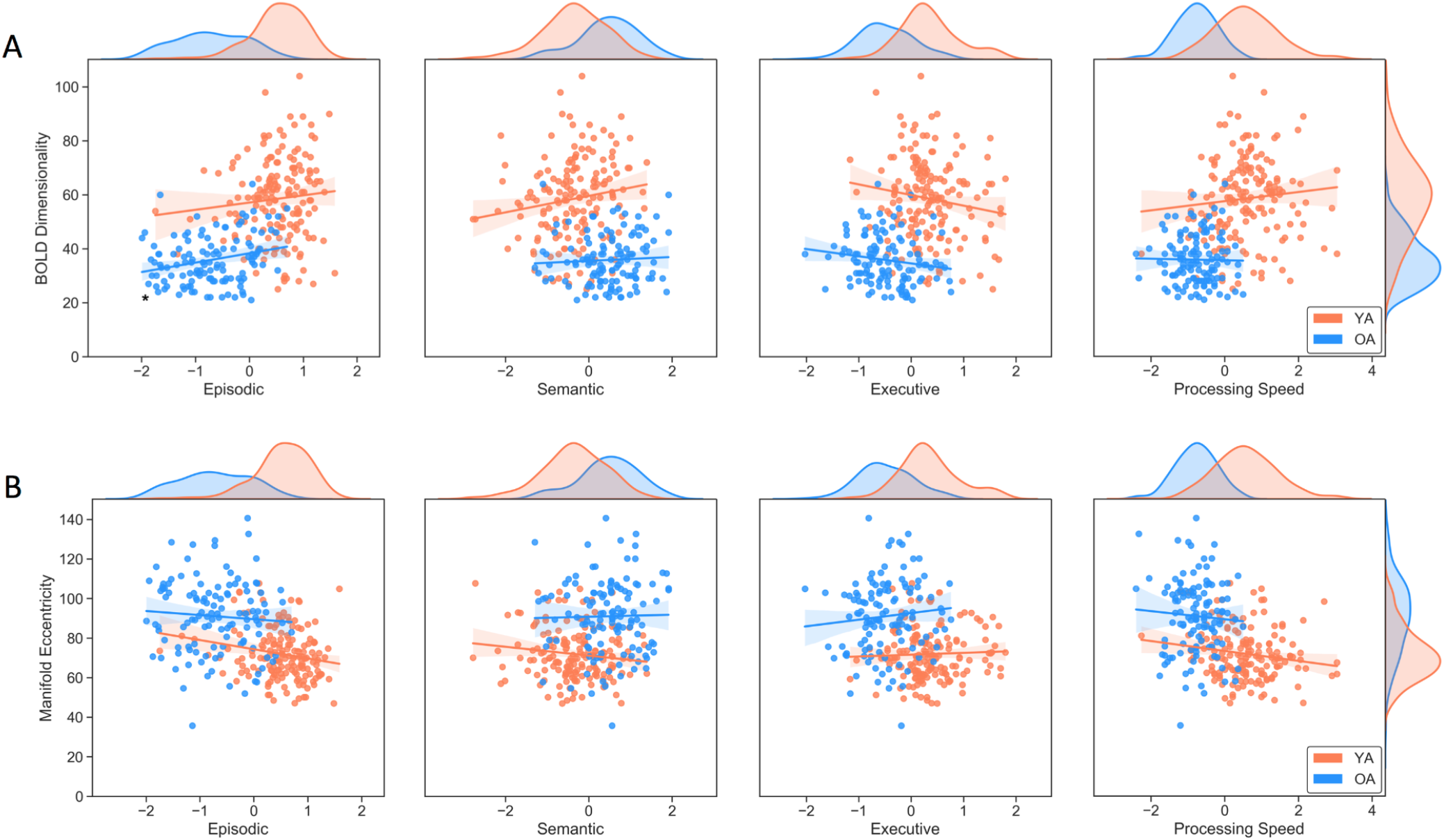
Scatterplots between cognitive scores and (A) BOLD dimensionality and (B) manifold eccentricity. Cognition distributions are shown at the top of each plot. Distributions for BOLD dimensionality and manifold eccentricity are shown in the rightmost plots. * indicates significant correlations. YA = younger adults; OA = older adults.

**Supplemental Figure 11:**
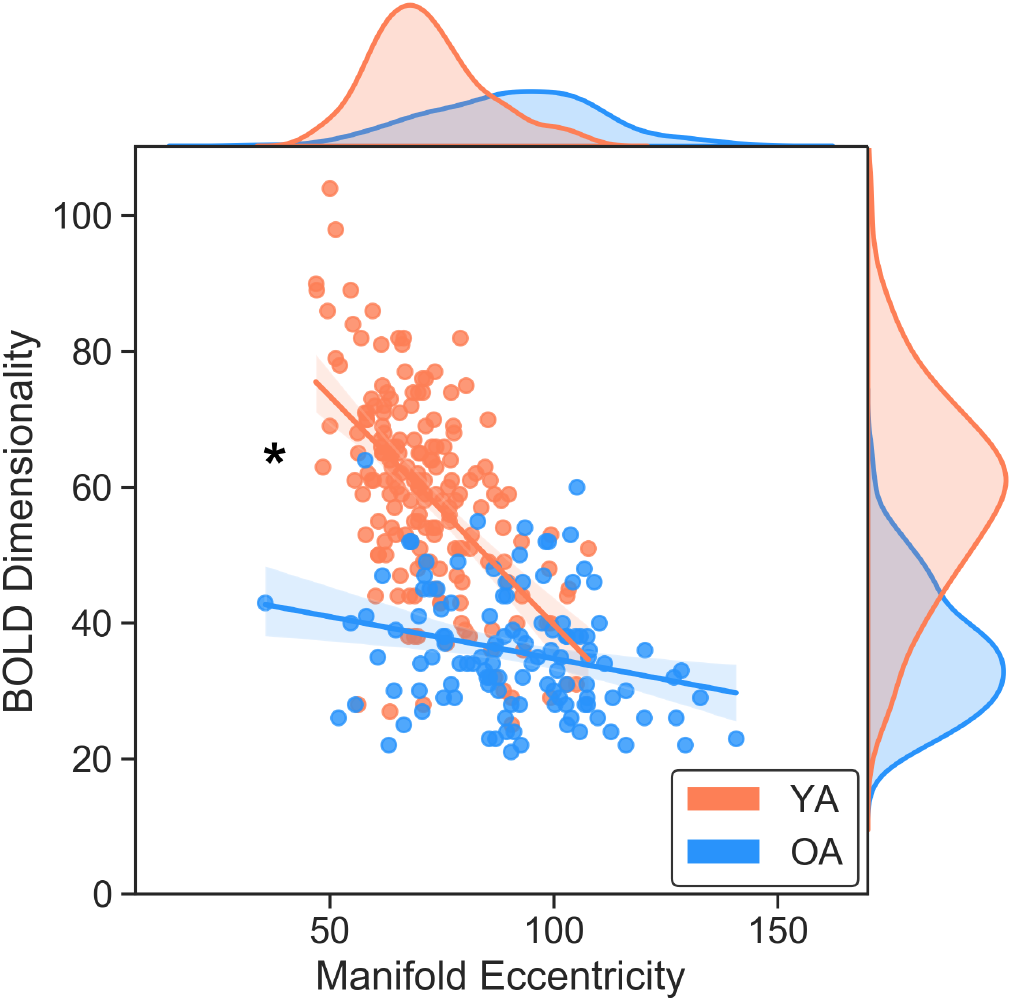
Scatterplot between BOLD dimensionality and manifold eccentricity. Distributions are shown on the respective axes. * indicates a significant difference between correlations. YA = younger adults; OA = older adults.

**Supplemental Figure 12:**
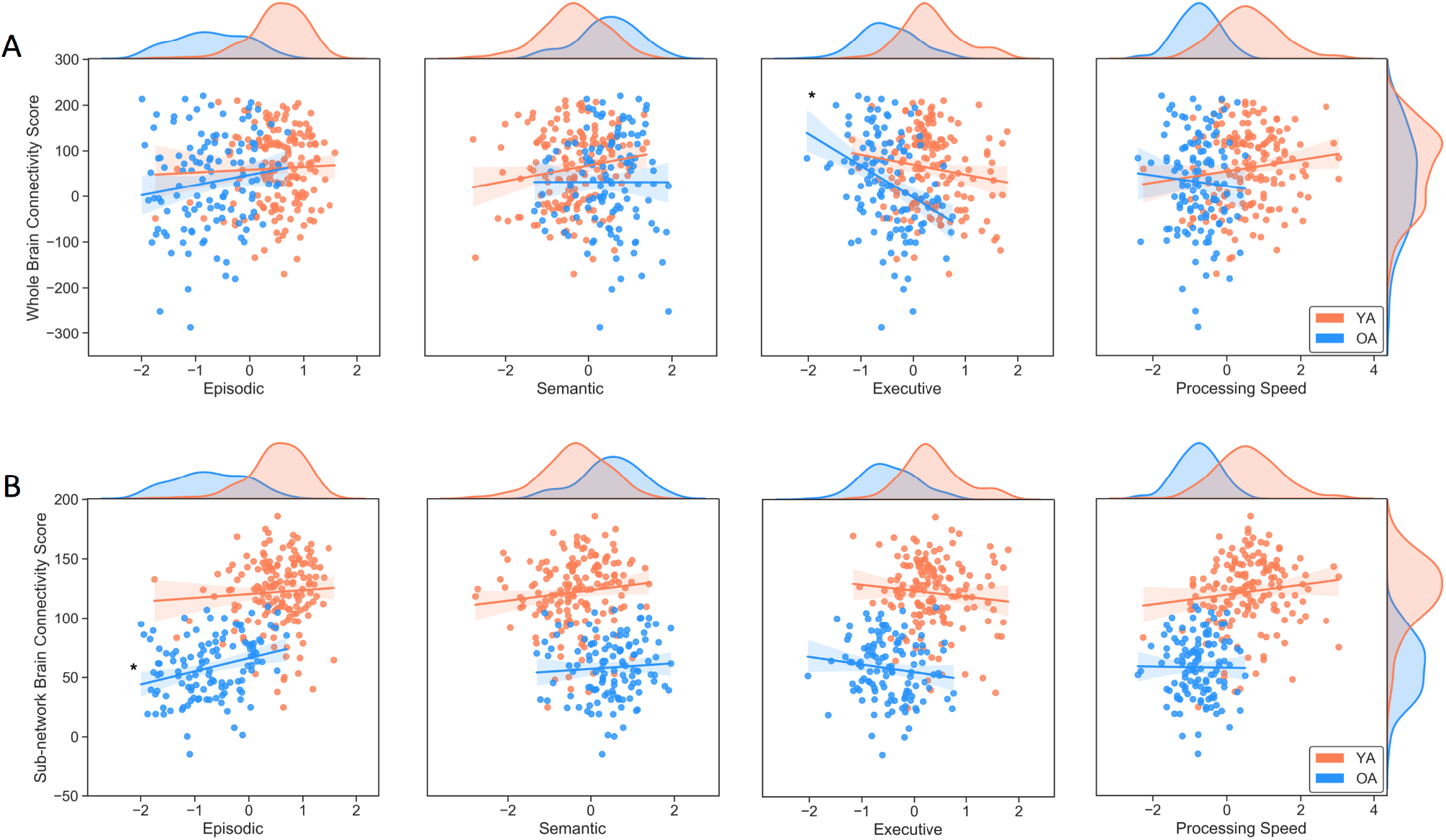
Scatterplots between cognitive scores and brain connectivity scores for the (A) full 7-network and (B) 3 sub-network analyses. Cognition distributions are shown at the top of each plot. Brain connectivity score distributions are shown in the rightmost plots. * indicates significant correlations. YA = younger adults; OA = older adults.

**Supplemental Figure 13:**
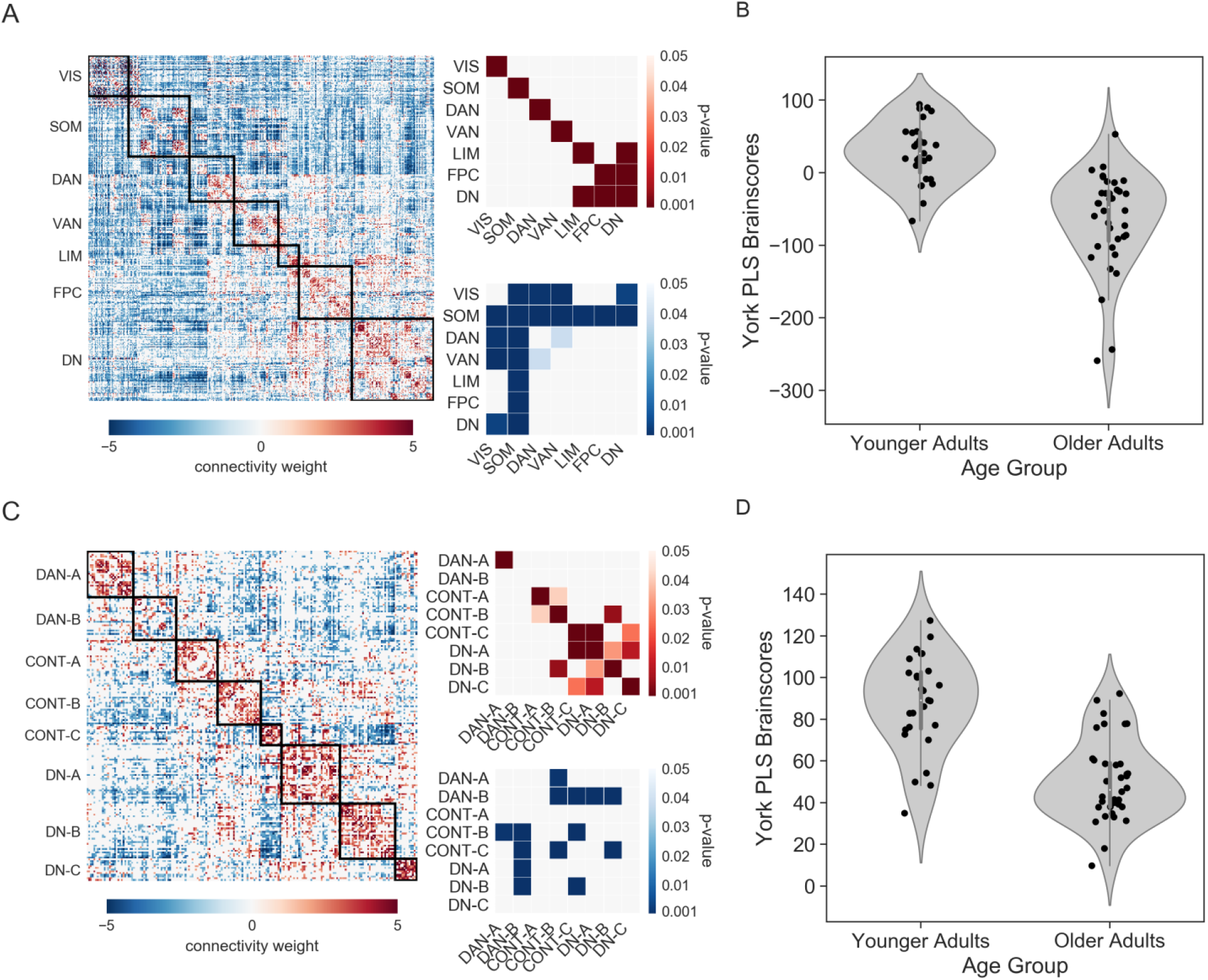
Replication of PLS results across sites. (A) Age differences in the full functional connectome for Ithaca participants. (B) Group differences in brain scores computed for York participants, based upon the edge-weights determined in the Ithaca sample. (C) Age differences in the functional connectivity of the default (DN), dorsal attention (DAN), and frontoparietal control (CONT) sub-networks from Ithaca participants. (D) Group differences in brain scores computed from Toronto participants, based upon the edge-weights from the sub-networks determined in Ithaca participants. DAN = dorsal attention, FPC = frontoparietal control, DN = default.

**Supplemental Table 2.** Brain-Behavior Correlations without Covariates.

| Factor | Younger Adults | Older Adults | Full Sample |
| --- | --- | --- | --- |
| **BOLD Dimensionality** |  |  |  |
| Episodic Memory | .094 (.235) [-.06, .24] | **.251 (.006) [.08, .41]** | .084 (.159) [-.03, .20] |
| Semantic Memory | .155 (.048) [.00, .03] | .057 (.539) [-.12, .23] | **.158 (.008) [.04, .27]** |
| Executive Function | -.146 (.062) [-.29, .01] | -.155 (.090) [-.33, .02] | **-.177 (.003) [-.29, -.06]** |
| Processing Speed | .096 (.221) [-.06, .25] | -.018 (.848) [-.20, .16] | .016 (.784) [-.10, .13] |
| **Manifold Eccentricity** |  |  |  |
| Episodic Memory | -.191 (.014) [-.34, -.04] | -.073 (.430) [-.25, .11] | -.084 (.157) [-.20, .03] |
| Semantic Memory | -.134 (.088) [-.28, .02] | .020 (.832) [-.16, .20] | -.077(.196) [-.19, .04] |
| Executive Function | .040 (.609) [-.11, .19] | .094 (.305) [-.09, .27] | .085 (.154) [-.03, .20] |
| Processing Speed | -.168 (.032) [-.31, -.01] | -.061 (.506) [-.24, .12] | -.080 (.182) [-.19, .04] |
| **Brain Connectivity Score (Whole Brain)** |  |  |  |
| Episodic Memory | .039 (.624) [-.12, .19] | .139 (.131) [-.04, .31] | .063 (.294) [-.05, .18] |
| Semantic Memory | .158 (.044) [.00, .30] | -.001 (.993) [-.18, .18] | .101 (.090) [-.02, .21] |
| Executive Function | -.145 (.064) [-.29, .01] | **-.350 (.000) [-.50, -.18]** | **-.258 (.000) [-.36, -.15]** |
| Processing Speed | .128 (.103) [-.03, .28] | -.062 (.504) [-.24, .12] | .027 (.647) [-.09, .14] |
| **Brain Connectivity Score (Sub-network)** |  |  |  |
| Episodic Memory | .059 (.452) [-.10, .21] | **.283 (.002) [.11, .44]** | .097 (.103) [-.02, .21] |
| Semantic Memory | .116 (.139) [-.04, .27] | .066 (.476) [-.11, .24] | .132 (.027) [.02, .24] |
| Executive Function | -.097 (.218) [-.25, .06] | -.132 (.150) [-.30, .05] | -.138 (.020) [-.25, -.02] |
| Processing Speed | .121 (.125) [-.03, .27] | -.011 (.908) [-.19, .17] | .023 (.697) [-.09, .14] |

*Note.* *r* values displayed with *p* values in parentheses and 95% confidence intervals in brackets. *pr* values displayed for the full sample with age as a covariate. Bold denotes significance with a Bonferroni correction at $p < .013$ for 4 tests.

**Supplemental Table 3.** Brain-Behavior Correlations with Covariates (Sex, Education, eWBV)

| Factor | Younger Adults | Older Adults | Full Sample |
| --- | --- | --- | --- |
| **BOLD Dimensionality** |  |  |  |
| Episodic Memory | .054 (.497) [-.10, .21] | **.243 (.007) [.07, .40]** | .056 (.351) [-.06, .17] |
| Semantic Memory | **.216 (.006) [.06, .36]** | .042 (.650) [-.14, .22] | **.174 (.003) [.06, .28]** |
| Executive Function | -.149 (.057) [-.30, .00] | -.154 (.094) [-.32, .03] | **-.182 (.002) [-.29, -.07]** |
| Processing Speed | .064 (.418) [-.09, .22] | -.011 (.903) [-.19, .17] | .009 (.884) [-.11, .13] |
| **Manifold Eccentricity** |  |  |  |
| Episodic Memory | -.185 (.018) [-.33, -.03] | -.092 (.319) [-.27, .09] | -.097 (.104) [-.21, .02] |
| Semantic Memory | -.184 (.019) [-.33, -.03] | .007 (.938) [-.17, .19] | -.094 (.114) [-.21, .02] |
| Executive Function | .046 (.557) [.11, .20] | .058 (.531) [-.12, .23] | .085 (.152) [-.03, .20] |
| Processing Speed | -.134 (.089) [-.28, .02] | -.101 (.272) [-.28, .08] | -.084 (.161) [-.20, .03] |
| **Brain Connectivity Score (Whole Brain)** |  |  |  |
| Episodic Memory | .068 (.390) [-.09, .22] | .056 (.543) [-.12, .23] | .006 (.919) [-.11, .12] |
| Semantic Memory | .126 (.108) [-.03, .27] | -.102 (.268) [-.28, .08] | .058 (.333) [-.06, .17] |
| Executive Function | -.136 (.084) [-.28, .02] | **-.320 (.000) [-.47, -.15]** | **-.249 (.000) [-.35, -.14]** |
| Processing Speed | .164 (.037) [.01, .31] | -.015 (.870) [-.19, .16] | .045 (.449) [-.07, .16] |
| **Brain Connectivity Score (Sub-network)** |  |  |  |
| Episodic Memory | .012 (.884) [-.14, .16] | **.229 (.012) [.05, .39]** | .031 (.602) [-.09, .15] |
| Semantic Memory | .169 (.031) [.02, .31] | .034 (.709) [-.15, .21] | .136 (.022) [.02, .25] |
| Executive Function | -.096 (.222) [-.25, .06] | -.145 (.115) [-.32, .04] | -.138 (.020) [-.25, -.02] |
| Processing Speed | .102 (.196) [-.05, .25] | -.016 (.862) [-.19, .16] | .019 (.753) [-.10, .14] |

*Note.* pr values displayed with *p* values in parentheses and 95% confidence intervals in brackets. Full sample includes age as a covariate. Bold denotes significance with a Bonferroni correction at $p < .013$ for 4 tests.

**Supplemental Table 4.** Brain-Behavior Correlations with Covariates (Sex, Education, eWBV, Site)

| Factor | Younger Adults | Older Adults | Full Sample |
| --- | --- | --- | --- |
| **BOLD Dimensionality** |  |  |  |
| Episodic Memory | -.094 (.232) [-.24, .06] | .182 (.047) [.00, .35] | -.010 (.864) [-.13, .11] |
| Semantic Memory | .126 (.109) [-.03, .27] | .076 (.409) [-.10, .25] | .129 (.030) [.01, .24] |
| Executive Function | -.008 (.916) [-.16, .15] | .126 (.171) [-.05, .30] | .013 (.823) [-.10, .13] |
| Processing Speed | -.059 (.454) [-.21, .10] | -.002 (.984) [-.18, .18] | -.023 (.703) [-.14, .09] |
| **Manifold Eccentricity** |  |  |  |
| Episodic Memory | -.082 (.296) [-.23, .07] | -.067 (.470) [-.24, .11] | -.061 (.306) [-.18, .06] |
| Semantic Memory | -.102 (.196) [-.25, .05] | -.001 (.988) [-.18, .18] | -.063 (.294) [-.18, .05] |
| Executive Function | -.088 (.264) [-.24, .07] | -.028 (.765) [-.21, .15] | -.034 (.574) [-.15, .08] |
| Processing Speed | -.042 (.599) [-.19, .11] | -.106 (.248) [-.28, .07] | -.069 (.250) [-.18, .05] |
| **Brain Connectivity Score (Whole Brain)** |  |  |  |
| Episodic Memory | -.096 (.222) [-.25, .06] | -.074 (.423) [-.25, .11] | -.101 (.091) [-.21, .02] |
| Semantic Memory | .007 (.929) [-.15, .16] | -.083 (.367) [-.26, .10] | -.024 (.683) [-.14, .09] |
| Executive Function | .026 (.740) [-.13, .18] | .030 (.749) [-.15, .21] | .017 (.778) [-.10, .13] |
| Processing Speed | .048 (.540) [-.11, .20] | .003 (.971) [-.18, .18] | .007 (.912) [-.11, .12] |
| **Brain Connectivity Score (Sub-network)** |  |  |  |
| Episodic Memory | -.130 (.098) [-.28, .02] | .161 (.079) [-.02, .33] | -.036 (.551) [-.15, .08] |
| Semantic Memory | .077 (.326) [-.08, .23] | .07 (.433) [-.11, .25] | .089 (.135) [-.03, .20] |
| Executive Function | .043 (.590) [-.11, .19] | .174 (.058) [-.01, .34] | .061 (.308) [-.06, .18] |
| Processing Speed | -.004 (.961) [-.16, .15] | -.002 (.981) [-.18, .18] | -.011 (.857) [-.13, .11] |

*Note.* *pr* values displayed with *p* values in parentheses and 95% confidence intervals in brackets. Full sample includes age as a covariate. Bold denotes significance with a Bonferroni correction at $p < .013$ for 4 tests.

**Supplemental Figure 14:**
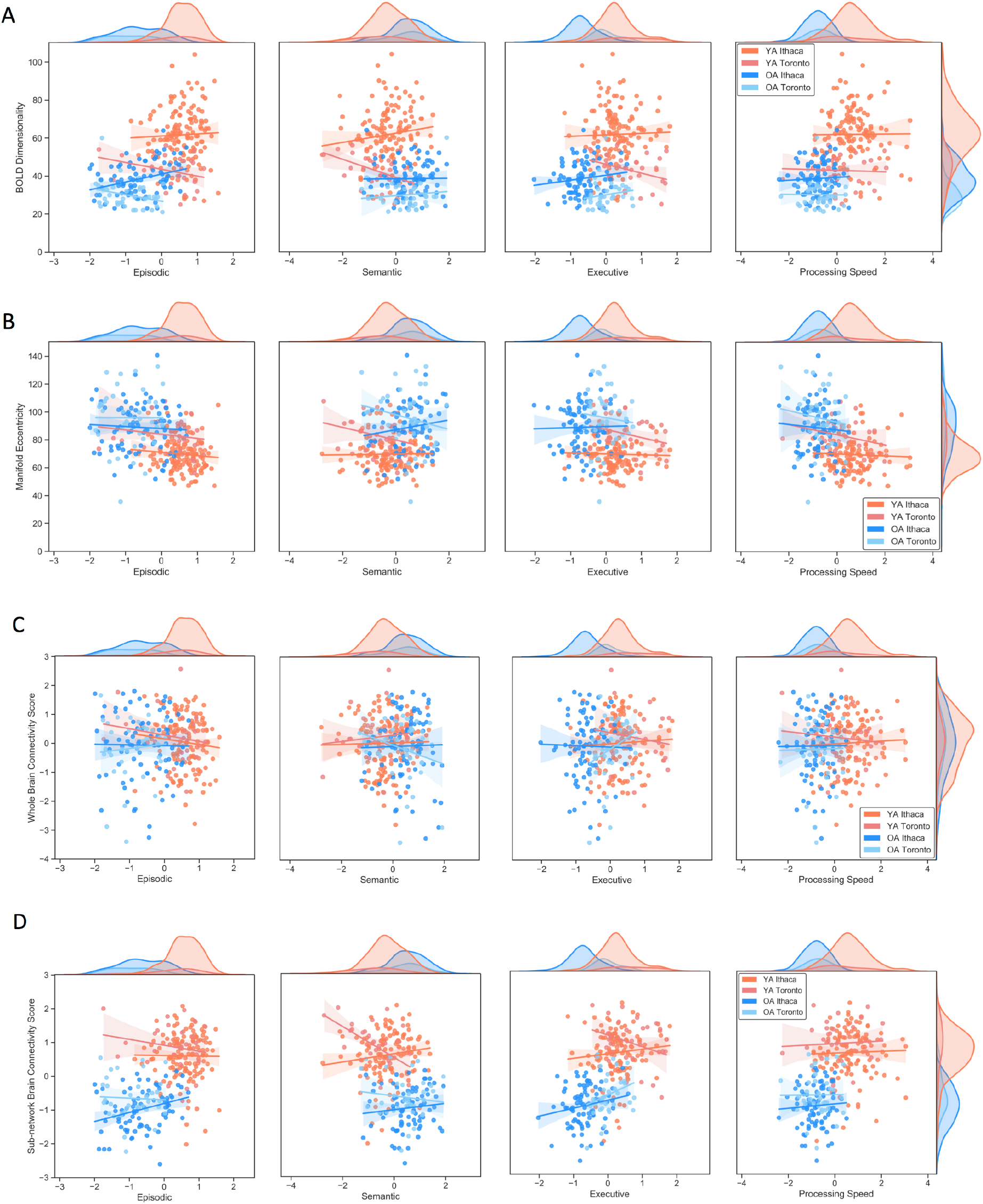
Scatterplots between cognitive scores and (A) BOLD dimensionality, (B) manifold eccentricity, (C) whole brain connectivity scores, and (D) sub-network brain connectivity scores by age group and site. Cognition distributions are shown at the top of each plot. Distributions for BOLD dimensionality, manifold eccentricity, and brain connectivity scores are shown in the rightmost plots. YA = younger adults; OA = older adults.

**Supplemental Table 5A.**
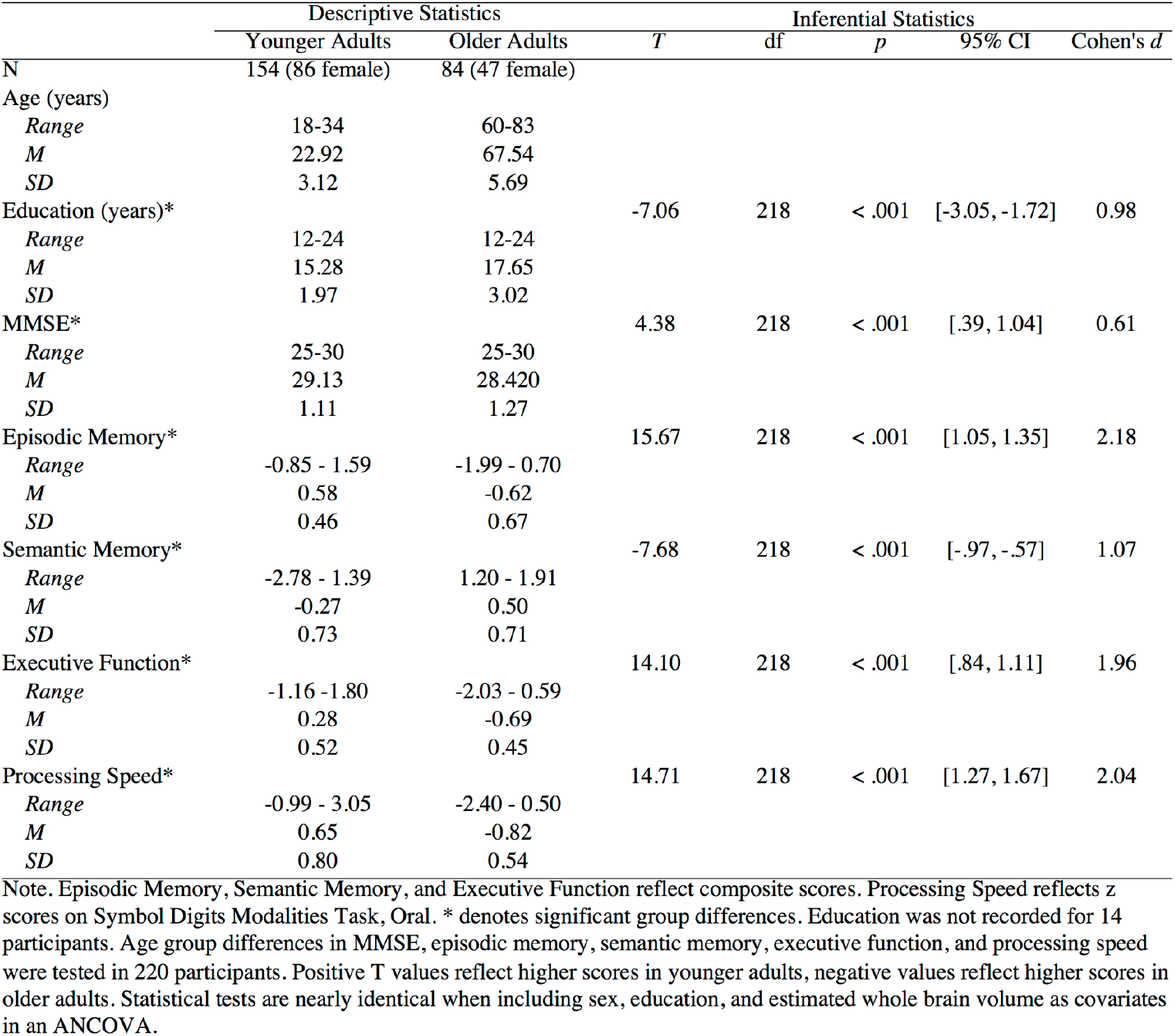
Sample Demographics: Ithaca.

**Supplemental Table 5B.**
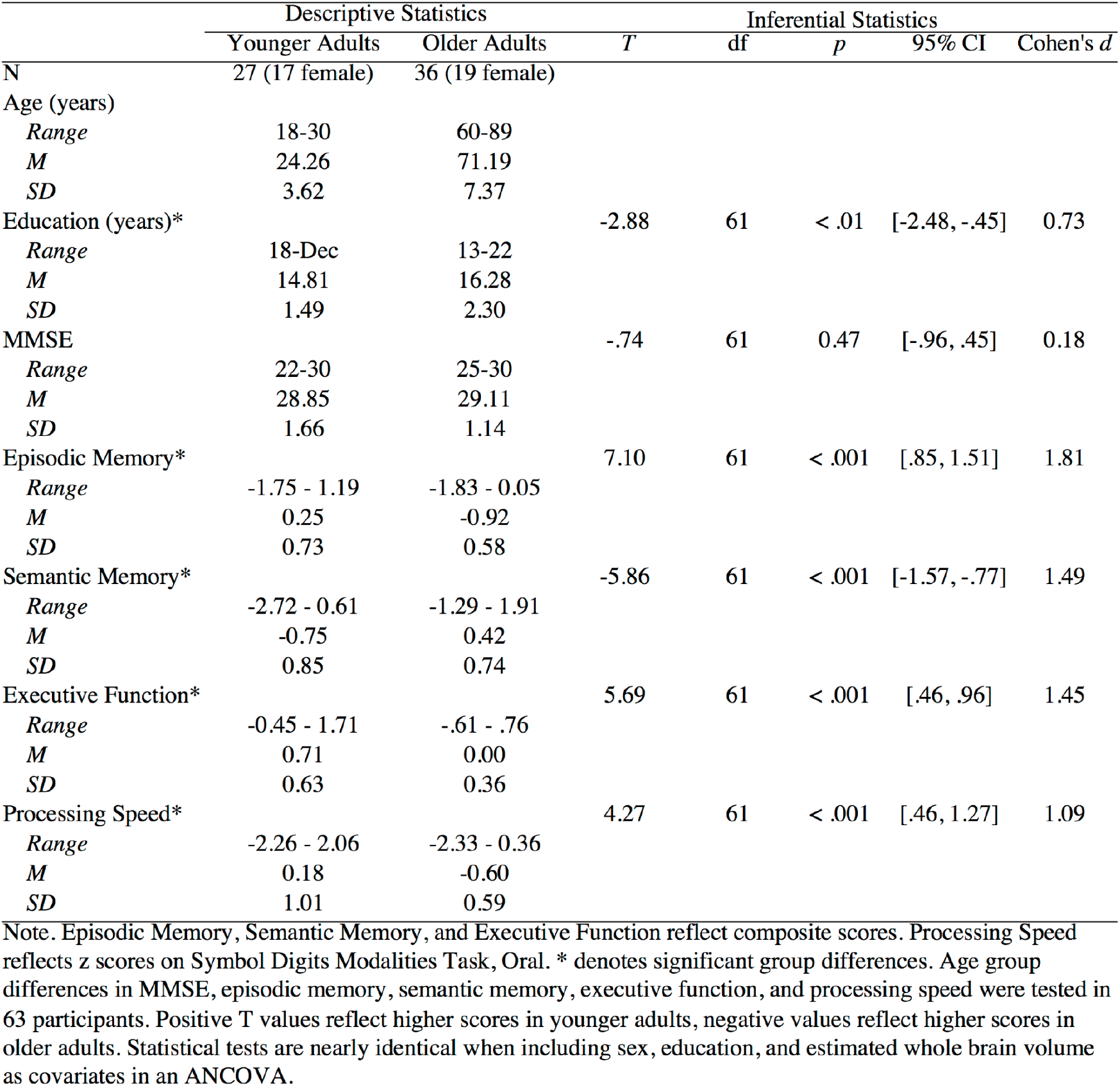
Sample Demographics: Toronto.

## Notes

### Competing Interest Statement

The authors have declared no competing interest.

